# Pre- and postsynaptic mechanisms of neuronal inhibition assessed through biochemically detailed modelling of GABAB receptor signalling

**DOI:** 10.1101/2025.03.19.644198

**Authors:** Tuomo Mäki-Marttunen, Jan Fredrik Kismul, Kadri Pajo, Jan Michael Schulz, Tiina Manninen, Gaute T. Einevoll, Marja-Leena Linne, Ole A. Andreassen, Jeanette Hellgren Kotaleski

## Abstract

GABAB receptors (GABABRs) are an important building block in neural activity. Despite their widely hypothesized role in many basic neuronal functions and mental disorder symptomatology, there is a lack of biophysically and biochemically detailed models of these receptors and the way they mediate neuronal inhibition. Here, we developed a computational model for the activation of GABABRs and its effects on the activation of G protein-coupled inwardly rectifying potassium (GIRK) channels as well as inhibition of voltage-gated Ca^2+^ channels. To ensure the generality of our modelling framework, we fit our model to electrophysiological data including patch-clamp and intracellular recordings that described both pre- and postsynaptic effects of the receptor activation. We validated our model using data on postsynaptic effects of GABABRs on layer V pyramidal cell firing activity ex vivo and in vivo and confirmed the strong impact of dendritic GIRK channel activation on the neuron output. Finally, we reproduced and dissected the effects of a knockout of RGS7 (a G protein signalling protein) on CA1 pyramidal cell electrophysiological properties, which shows the potential of our model in generating insights on genetic manipulations of the GABABR system and related genetic variants. Our model thus provides a flexible tool for biochemically and biophysically detailed simulations of different aspects of GABABR activation that can reveal both foundational principles of neuronal dynamics and brain disorder-associated traits and treatment options.

## 1 Introduction

Computational models of neurons and neuronal networks have emerged as a useful tool to obtain knowledge about the central nervous system and how its functions arise as a complex interplay of various neurochemical processes. *γ*-aminobutyric acid (GABA) receptors of type B (GABABRs) represent one essential component of neural activity in both cortical and subcortical brain regions [Bettler et al., 2004, Groen, 2011, Tureček et al., 2023]. In contrast to the better understood and more extensively targeted ionotropic GABA receptors of type A (GABAARs) [Ghit et al., 2021], GABABRs are G-protein coupled receptors and thus the inhibition they exert is slow and dependent on a larger intracellular cascade [Bettler et al., 2004]. Activation of the GABABR releases the beta-gamma subunit of the G_i_ protein (G_i*βγ*_) that, upon binding to ion channels, inhibits N-type voltage-gated Ca^2+^ channels (VGCCs; typically presynaptic) and activates G protein-coupled inwardly rectifying K^+^ (GIRK) channels (typically postsynaptic) [Kohl and Paulsen, 2010]. In addition, GABABRs have been shown to directly inhibit postsynaptic L-type Ca^2+^ channels in some neuron types [Pérez-Garci et al., 2006, Pérez-Garci et al., 2013, Schulz et al., 2021]. The impact of GABABR activation is complex as it also mediates short-term synaptic depression in both excitatory and inhibitory presynaptic terminals [Isaacson et al., 1993, Olpe et al., 1994]. This presents a challenge for mechanistic analysis of GABABR-mediated effects in single neurons as well as neural circuits: namely, the effects of manipulations of the GABABR system are difficult to predict and the observed phenomena are difficult to generalize between neuron types [Craig and McBain, 2014]. Overcoming this challenge is important due to the association of the GABABR pathway with heritable mental disorders such as schizophrenia [Trubetskoy et al., 2022] and its promise as a drug target [Evenseth et al., 2020]. Here, we propose the use of biochemically detailed computational modeling to enhance our understanding of GABABR-mediated effects and to help address this challenge.

Traditionally, computational models of neurons and neuronal networks have heavily relied on GABAAR-mediated currents, often excluding GABABR-mediated currents or simplifying them as static, long-tailed inhibitory currents [Traub et al., 1994, Turi et al., 2019]. While models of GABABR-mediated inhibition that describe some of the intracellular machinery exist, they only focus on either post[Destexhe, 1998, Park et al., 2014] or presynaptic [Li et al., 2020] effects, and they do not include the contribution of key proteins involved such as regulators of G protein signalling (RGS) proteins. RGS proteins inhibit G-protein activity by hydrolysing the G_i*α*_-bound GTP to GDP, permitting the G_i*α*_ subunit to reassociate with G_i*βγ*_ to inactivate the G_i_ protein [Zhong et al., 2003]. Here, we aimed to develop a detailed modeling framework to enable investigations of novel GABABR features not included in current models. We fit a mass-action-law-based model including GABABRs, G_i_, RGS proteins and GIRK and N-type Ca^2+^ channels to electrophysiological patch-clamp and intracellular recording data [Isaacson et al., 1993, Olpe et al., 1994] and present a biophysically detailed modelling framework in which both pre- and postsynaptic effects of GABABR manipulations can be quantified. We verified the appropriate functioning of the presynaptic depression model using a detailed model of presynaptic terminal integrated into an axon segment and incorporated a neurotransmitter release model. We then validated the postsynaptic model using additional data on effects of GABABR activation on layer 5 pyramidal cell (L5PC) firing under patch-clamp stimulus [Schulz et al., 2021] or electrical paw stimulation [Palmer et al., 2012] and explored the relationship between the effects of pre- and postsynaptic GABABR activation on L5PC firing. We employed our model to dissect the causes of altered excitability and plasticity in CA1 pyramidal cells of RGS7-KO mice [Ostrovskaya et al., 2014]. This shows that our model can be used for modelling of genetic manipulations of GABABR pathway and is thus immediately useful for exploring mechanisms and possible treatments of mental disorders.

## 2 Methods

In this work, we first developed a single-compartment model of GABAB receptor activation without incorporating membrane ion-channel mechanisms. However, to demonstrate its potential in neuroscience research, we integrated it into various Hodgkin-Huxley-type models. Table 1 summarizes the models employed in this work.

**Table 1:** Summary of models. The models developed in this work (*,†,‡) were based on existing models: [Zhong et al., 2003, Jedrzejewska-Szmek et al., 2017, Mäki-Marttunen et al., 2020]*, [Destexhe et al., 1994, Migliore et al., 1995, Hay et al., 2011]†, [Wang, 1999]‡.

| Model | Number of compartments | References | Where described | Where employed |
| --- | --- | --- | --- | --- |
| Single-compartment GABABR activation model | 1 or more | (developed here *) | Sec. 2.1 | Figs. 1–5 |
| Model of an axon and axon terminal (presynaptic) | 5 | (developed here †) | Sec. 2.2 | Figs. 2, 4 |
| Layer V pyramidal cell model (postsynaptic) | 196 | [Hay et al., 2011] | Sec. 2.3 | Figs. 3, 4 |
| CA1 pyramidal cell model (postsynaptic) | 144 | [Combe et al., 2018] | Sec. 2.4 | Fig. 5 |
| Neurotransmitter release model (cross-synaptic) | 1 or more | (developed here ‡) | Sec. 2.5 | Fig. 4 |

### 2.1 Construction of the GABABR activation model

#### 2.1.1 Modelling intracellular signalling networks underlying G_i*βγ*_ activation and their effects on ion channels

We modeled the intracellular signalling in the GABABR-containing subcellular compartments (presynaptic terminal, post-synaptic spine, or a dendritic compartment) using the law of mass action. We only used deterministic simulations. The reactions as well as the default initial concentrations and reaction rates were adopted from [Jedrzejewska-Szmek et al., 2017, Mäki-Marttunen et al., 2020] (apart from the RGS reactions that were taken from [Zhong et al., 2003]) and are listed in Table 2A.

**Table 2:** The biochemically detailed model of GABABR activation. **A**: Reactions describing the activation of GABABRs and G_i_ protein and the binding of G_i*βγ*_ to GIRK and N-type VGCCs. The reaction rates were adopted from [Jedrzejewska-Szmek et al., 2017, Mäki-Marttunen et al., 2020, Zhong et al., 2003]: some reaction rates were used as such, while others were multiplied by coefficients *k*_1_, *k*_2_, *k*_3_, *k*_4_, and *k*_5_ that were varied in a multiobjective optimisation framework (see (B)). The unit of *k*_*b*_ is always 1/ms and the unit of *k*_*f*_ is 1/ms or 1/(nM·ms) when the reaction has one or two reactants, respectively. **B**: The ranges for all optimized parameters. The parameters *k*_1_, *k*_2_, *k*_3_, *k*_4_, and *k*_5_ are coefficients for the reaction rates of reactions R1–R7 and R11–R15 (see (A)). The parameter *k*_2*x*_ was determined so that the affinity was 110 nM as measured experimentally [Lehmann et al., 2009], i.e., *k*_2*x*_ = 5.555 · 10^*-*6^/(nM · ms) *× k*_2_ *×* 110 nM. In addition, the initial concentration of the RGS proteins in all domains and the GABA flux were varied. The concentrations of the rest of the species were [GABABR] = 0.4 µM, [G_i*βγ*_] = 2.6 µM, [GIRK] = 1 µM, and [VGCC] = 0.1 µM, unless otherwise stated. Both the reactions (A) and initial concentrations (B) were the same in pre- and postynaptic compartments, apart from those of VGCCs (R15), which were absent from postsynaptic compartments, GIRK channels (R11-R14), which were absent from presynaptic compartments, and RGS proteins (R8-R9), whose concentration differed between presynaptic terminals of E*→* E and I*→* E synapses and postsynaptic compartments (B). The zero concentration of GIRK channels effectively blocks reactions R11–R14 at the presynaptic terminals while the zero concentration of VGCCs in effect blocks reaction R15 at the postsynaptic compartments.

| **A** |  |  | **B** |  |  |
| --- | --- | --- | --- | --- | --- |
| Reactions | $k_f$ | $k_b$ | Parameter | Lower limit | Upper limit |
| R1: gaba → gabaOut | $k_1 \times 0.0005$ | 0.0 | $k_1$ | 0.0 | 1000 |
| R2: gaba + GABABR ↔ gabaGABABR | $k_2 \times 5.555 \cdot 10^{-6}$ | $k_{2x}$ | $k_2$ | 0.0001 | 1000 |
| R3: gaba + GABABRG<sub>i</sub> ↔ gabaGABABRG<sub>i</sub> | $k_2 \times 5.555 \cdot 10^{-6}$ | $k_{2x}$ | $k_3$ | 0.0 | 1000 |
| R4: GABABR + G<sub>i</sub> ↔ GABABRG<sub>i</sub> | $k_3 \times 7.5 \cdot 10^{-5}$ | $k_3 \times 0.000125$ | $k_4$ | 0.0 | 100 |
| R5: gabaGABABR + G<sub>i</sub> ↔ gabaGABABRG<sub>i</sub> | $k_3 \times 0.00015$ | $k_3 \times 0.00025$ | $k_5$ | 0.0 | 100 |
| R6: gabaGABABRG<sub>i</sub> → gabaGABABRG<sub>i\beta\gamma</sub> + GiaGTP | $k_3 \times 0.000125$ | 0.0 | [RGS<sub>post</sub>] | 0.1 $\mu\text{M}$ | 10 $\mu\text{M}$ |
| R7: gabaGABABRG<sub>i\beta\gamma</sub> → gabaGABABR + G<sub>i\beta\gamma</sub> | $k_3 \times 0.001$ | 0.0 | [RGS<sub>pre,E</sub>] | 0.1 $\mu\text{M}$ | 10 $\mu\text{M}$ |
| R8: GiaGTP + RGS ↔ GiaGTPRGS | $2 \cdot 10^{-6}$ | 0.002 | [RGS<sub>pre,I</sub>] | 0.1 $\mu\text{M}$ | 10 $\mu\text{M}$ |
| R9: GiaGTPRGS → GiaGDP + RGS | 0.03 | 0.0 | $f_{\text{GABA}}$ | 0 | 3000/ms |
| R10: GiaGDP + G<sub>i\beta\gamma</sub> → G<sub>i</sub> | 0.00125 | 0.0 | | | |
| R11: GIRK + G<sub>i\beta\gamma</sub> ↔ GIRKG<sub>i\beta\gamma</sub> | $k_4 \times 1.4 \cdot 10^{-5}$ | $k_5 \times 0.001$ | | | |
| R12: GIRKG<sub>i\beta\gamma</sub> + G<sub>i\beta\gamma</sub> ↔ GIRKG<sub>i\beta\gamma</sub>2 | $k_4 \times 1.4 \cdot 10^{-5}$ | $k_5 \times 0.001$ | | | |
| R13: GIRKG<sub>i\beta\gamma</sub>2 + G<sub>i\beta\gamma</sub> ↔ GIRKG<sub>i\beta\gamma</sub>3 | $k_4 \times 1.4 \cdot 10^{-5}$ | $k_5 \times 0.001$ | | | |
| R14: GIRKG<sub>i\beta\gamma</sub>3 + G<sub>i\beta\gamma</sub> ↔ GIRKG<sub>i\beta\gamma</sub>4 | $k_4 \times 1.4 \cdot 10^{-5}$ | $k_5 \times 0.001$ | | | |
| R15: VGCC + G<sub>i\beta\gamma</sub> ↔ VGCCG<sub>i\beta\gamma</sub> | $k_4 \times 1.4 \cdot 10^{-5}$ | $k_5 \times 0.001$ | | | |

The binding of G_i*βγ*_ to GIRK (postsynaptic) or N-type Ca^2+^ (presynaptic) channels was considered as the main output of the model. The GIRK channel was activated in a dose-dependent manner by binding of G_i*βγ*_: the conductance was 0 when non-bound and 1%, 6%, 26%, or 100% of the maximal conductance when bound by 1, 2, 3, or 4 G_i*βγ*_ molecules [Dascal and Kahanovitch, 2015]. The VGCCs, in turn, was inhibited (only one G_i*βγ*_ molecule per channel was required for this) such that the activation curve was shifted by +10 mV and the activation time constant was made 30% larger in the G_i*βγ*_-bound compared to the non-bound VGCCs [Huynh et al., 2015].

#### 2.1.2 Model fitting

To fit the rate coefficients governing the GABABR activation dynamics, we used a multiobjective optimization algorithm, NSGA-II [Deb et al., 2002], as implemented in [Bahl et al., 2012, Mäki-Marttunen et al., 2018]. We simultaneously adjusted the model of Table 2 to fit three data sets: 1) GABABR-mediated inhibitory postsynaptic currents (IPSCs) [Isaacson et al., 1993], 2) GABABR-mediated short-term synaptic depression of excitatory-to-excitatory (E*→*E) synapses [Isaacson et al., 1993], and 3) GABABR-mediated short-term synaptic depression of inhibitory-to-excitatory (I*→*E) synapses [Olpe et al., 1994]. All three data sets were obtained by Schaffer collateral stimulation and intracellular recordings from CA1 pyramidal cells [Isaacson et al., 1993, Olpe et al., 1994]. We varied the reaction rates for GABA uptake from the extracellular medium (*k*_1_), the forward rate of GABA binding to the GABABR (*k*_2_), a multi-reaction rate coefficient for G_i_ protein activation (*k*_3_), and the forward (*k*_4_) and backward (*k*_5_) reaction rates for G_i*βγ*_ binding to the voltage-gated ion channels, namely, GIRK and N-type Ca^2+^ channel (see Table 2B). We also varied the magnitude of the GABA flux into the vicinity of the synapse and the RGS concentrations. All varied parameters applied to all three types of simulation, except for the RGS concentration, which was allowed to vary between the synapse types (postsynaptic E*→*E, presynaptic E*→*E, presynaptic I*→*E) to capture the differences in the dynamics of GABABR effects [Isaacson et al., 1993, Olpe et al., 1994].

We introduced two objective functions to constrain the model responses according to the electrophysiological data. The first objective function quantified the difference between predicted postsynaptic GIRK conductance and the GABABR-mediated IPSCs measured in [Isaacson et al., 1993], both normalized by their maximal values:

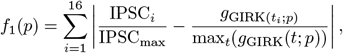

where *t*_*i*_, i=1,…,16 are the time instants of the IPSC measurements and IPSC_*i*_ are the corresponding IPSC values. Here, *g*_GIRK_(*t*; *p*) is the predicted GIRK conductance at time *t* after stimulus onset given model parameters *p*. It was determined from the simulation of postsynaptic sections (no VGCCs, 1 µM GIRK channels, which corresponds to 120 molecules/µm^2^ in a compartment with a diameter of 0.8 µm, and RGS concentration set according to the parameter given for postsynaptic sections) using the predicted concentrations of GIRK channels in different states as

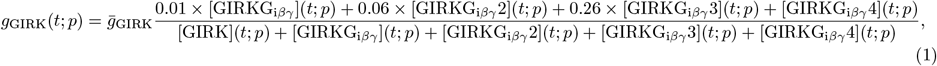

where 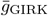 is the theoretical maximum (here, we used the value 33 pS [Chen and Johnston, 2005]) for GIRK channel conductance and [GIRKG_i*βγ*_**X**](t;p), **X**=1..4 is the predicted concentration of GIRK channels bound with **X** G_i*βγ*_ molecules at time *t*, given parameters *p*. The second objective function quantified the difference between predicted VGCC binding and the experimentally observed paired-pulse ratios (PPR), both normalized, across the two presynaptic types:

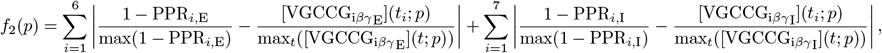

where PPR_*i*,E_ and PPR_*i*,I_ are the PPRs in E*→*E or I*→*E synapses as measured in [Isaacson et al., 1993] or [Olpe et al., 1994], respectively, for the *i*th time point *t*_*i*_. Here, [VGCCG_i*βγ*_E](*t*; *p*) and [VGCCG_i*βγ*_I](*t*; *p*) are the predicted concentration of G_i*βγ*_-bound VGCC in E*→*E or I*→*E synapses, respectively, at time *t*, given parameters *p*. We also introduced a third objective function that maximizes the absolute signal amplitudes in the three types of simulations:

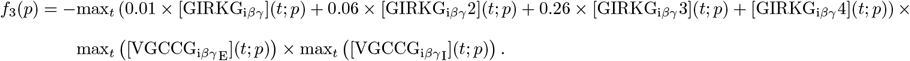

We ran the NSGA-II optimization algorithm with 2000 sample parameter sets for 25 generations. The third objective function was not used for selection of the final parameter set but as a signal to avoid obtaining models that produce correct G_i*βγ*_ binding dynamics when normalized but little total G_i*βγ*_ binding. We picked the parameter set that performed best in the two first objectives compared to median error values, i.e. *p*_final_ = argmin*p* (*f*_1_(*p*)*/*median_*p*_ (*f*_1_(*p*)) + *f*_2_(*p*)*/*median_*p*_ (*f*_2_(*p*))).

### 2.2 Presynaptic modelling: Composite model of an axon and a GABABR-containing presynaptic terminal

In the simulations of GABABR-mediated short-term synaptic depression, we modelled the presynaptic terminal (axonal bouton) surrounded by axonal compartments. The axonal bouton was modelled as a single compartment cylinder with 0.8 µm length and 0.4 µm diameter located between two active axon segments with 20 µm length and 1 µm diameter. On both ends of the active axon there were 100 µm long, 1 µm wide passive axon segments. For the active segments, we implemented the active membrane mechanisms and the Ca^2+^ decay mechanism from the active-axon model of [Hay et al., 2011]. For the axonal bouton, we adapted the N-type Ca^2+^ channel model from [Migliore et al., 1995] by adding the description of the “reluctant” mode of activity, which simulates the channel behaviour under G_i*βγ*_-binding. This was implemented such that a fraction (determined by the intracellular signalling model as 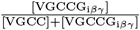) of the channels were in the reluctant mode while the rest 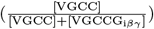 remained in the “willing” mode. The activation midpoint was 10 mV higher and the time constant was 31% larger in the reluctant than in the willing mode [Colecraft et al., 2000, Huynh et al., 2015]. We assumed the passive axon segments to be completely myelinated and thus void of any transmembrane currents.

We stimulated the end of the (passive) axon by a square-pulse current with amplitude 0.3 nA and 0.5 ms duration to induce an action potential (AP) in the active axon compartments and the axonal bouton. The AP caused a Ca^2+^ influx through the N-type Ca^2+^ channels. We introduced the model of Ca^2+^-induced neurotransmitter release from [Destexhe et al., 1994]. Namely, we introduced the reactions involving Ca^2+^ binding to a fusion factor molecule and further to a neurotransmitter-containing vesicle to induce the release in the axonal bouton (in addition to the GABABR model) — see Table 3.

**Table 3:** Table of reactions and initial concentrations between molecular species involved in Ca^2+^-induced release of neurotransmitter.

| Reactions | $k_f$ | $k_b$ | Initial concentrations |
| --- | --- | --- | --- |
| R16: $4 \times \text{Ca} + \text{Ffactor} \rightleftharpoons \text{FfactorCa4}$ | $50 \times 10^9 \frac{1}{\text{ms} \cdot \text{mM}^4}$ | $0.1 \frac{1}{\text{ms}}$ | $[\text{Ffactor}] = 1 \mu\text{M}$ |
| R17: $\text{FfactorCa4} + \text{Ves} \rightleftharpoons \text{FfactorCa4Ves}$ | $100000 \frac{1}{\text{ms} \cdot \text{mM}}$ | $0.1 \frac{1}{\text{ms}}$ | $[\text{Ves}] = 1 \mu\text{M}$ |
| R18: $\text{FfactorCa4Ves} \rightarrow 10000 \cdot \text{NT} + \text{Ffactor} + \text{Ves} + 4 \cdot \text{Ca}$ | $4 \frac{1}{\text{ms}}$ | 0 | Other species: $0 \mu\text{M}$ |
| R19: $\text{NT} \rightarrow \{\}$ | $10 \frac{1}{\text{ms}}$ | 0 | |

### 2.3 Postsynaptic modelling: Layer 5 pyramidal cell model and inclusion of GIRK channel electrophysiology

We used the Hay model of thick-tufted L5PCs [Hay et al., 2011] to analyze the postsynaptic effects of GABABR activation. The model includes Hodgkin-Huxley-type description of 11 ion-channel mechanisms (namely, the passive leak, transient and persistent Na^+^, transient and persistent K^+^, Kv3.1- and M-type K^+^, SK, HCN, and high- and low-voltage-activated Ca^2+^ currents) and a description of intracellular Ca^2+^ ion dynamics [Hay et al., 2011]. We introduced the GABABR activation model (Table 2) in all of the dendritic sections using the NEURON RxD formalism [McDougal et al., 2013]. We then introduced a GIRK current model based on an existing Kir channel model [Stegen et al., 2012, Yim et al., 2015] in these sections and coupled the conductance of this channel to the intracellular signalling model. In the experiments of Section 3.3, all dendritic sections contained the GABABR activation model and the interacting GIRK channels, whereas in the experiments of Section 3.4, only those dendritic sections that received synaptic inputs included the GABABR and GIRK models. To model the bath application of the agonist, we introduced an additional backward reaction GABA *←* GABA_out_ with the same rate as the forward reaction R1 (see Table 2A). In this model, the GIRK current was described as

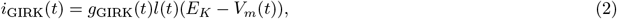

where *g*_GIRK_(*t*) is as in Eq. 1, *l*(*t*) is a gating variable, *E*_*K*_ is the reversal potential of K^+^, and *V*_*m*_ is the membrane potential. The gating variable obeys the differential equation

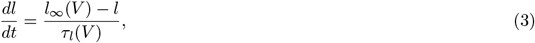

where 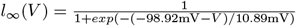 and 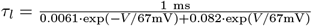.

### 2.4 Postsynaptic modelling: CA1 pyramidal cell model and inclusion of GIRK channel electrophysiology

We used the Combe model of CA1 pyramidal cells [Combe et al., 2018] to analyze the postsynaptic GABABR activation in CA1 pyramidal cells and its effects on the cell’s excitability and response to plasticity-inducing stimulation. The model includes Hodgkin-Huxley-type description of 12 ion-channel mechanisms (the passive leak, transient and persistent Na^+^, delayed rectifier K^+^, A- and M-type K^+^, slow- and medium AHP (Ca^2+^-activated) K^+^, L-, R- and T-type Ca^2+^, and HCN currents) and a description of intracellular Ca^2+^ ion dynamics [Combe et al., 2018, Bianchi et al., 2012]. Similar to the L5PCs, we introduced the GABABR activation model (Table 2) in all apical and basal dendritic sections together with the GIRK current model (Eq. 2). When determining the Ca^2+^ currents through the N-methyl-D-aspartate (NMDA) receptors, we multiplied the total NMDAR-conducted currents by 0.1 as approximately 10% of NMDAR-conducted were estimated to be Ca^2+^ currents [Burnashev et al., 1995].

### 2.5 Cross-synaptic modelling: Neurotransmitter-gated model of AMPAR- and NMDAR-mediated currents

For the simulations assessing the effects of presynaptic GABABRs on postsynaptic neuron activity, we integrated the neurotransmitter release model (Table 3), regulated by the GABABR activation model (Table 2), into an existing synaptic model. Namely, we adapted the *α*-amino-3-hydroxy-5-methyl-4-isoxazolepropionic acid (AMPA)/NMDA synapses of [Wang, 1999, Mäki-Marttunen and Mäki-Marttunen, 2022] to make them activated by the release of neurotransmitter in the axonal bouton in a dose-dependent manner. The AMPAR conductance was described by *g*_AMPA_ = *g*_AMPA,max_*s*_AMPA_ where *s*_AMPA_ was determined by

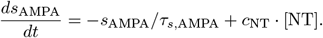

Here, *τ*_*s*,AMPA_ = 2 ms is the time constant of the decay of the AMPAR-mediated current, [NT] is the concentration of the neurotransmitter (glutamate) in the synaptic cleft and *c*_NT_ (unit 1/ms/mM) is the constant relating the neurotransmitter concentration to the steepness of the rise of the AMPAR conductance. Similarly, the NMDAR conductance was described by *g*_NMDA_ = *g*_NMDA,max_*s*_NMDA_ where *s*_NMDA_ was determined by

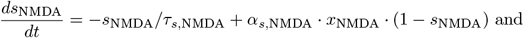

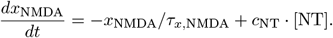

Here, *τ*_*x*,NMDA_ = 2 ms and *τ*_*s*,NMDA_ = 100 ms are the time constants of the rise and decay, respectively, of the NMDAR-mediated current, and the neurotransmitter gating is mediated by the coupling of [NT] and *x*_NMDA_ in a similar fashion as the coupling of [NT] and *s*_AMPA_.

### 2.6 Code availability

All simulations were run using NEURON v. 8.2.6, using Python (3.9.20) interface. Our simulation scripts are available at ModelDB (accession number 2018268, password ‘gabab’ required during peer review). To ensure easier model re-use and accordance with the FAIR data principles [Wilkinson et al., 2016], we also provide the single-compartment GABABR biochemical signalling model in SBtab and SBML formats. The model with three experimental settings was manually transferred into human-readable spreadsheet-based SBtab format [Lubitz et al., 2016] from which it was converted into an SBML file via the SBtabVFGEN tool as part of the modeling workflow by [Santos et al., 2022].

## 3 Results

### 3.1 The model fits experimental data on presynaptic short-term depression and postsynaptic GIRK currents

To obtain a realistic and generalizable model of GABABR activation leading to GIRK channel activation and N-type Ca^2+^ channel inactivation, we fit the reaction rates to electrophysiological data relating to both pre- and postsynaptic phenomena. The time course of postsynaptic GIRK channel activity has been measured in CA1 pyramidal cells [Isaacson et al., 1993], where the GABABR-mediated GIRK currents peaked at 200 ms and returned to zero in approximately 800 ms. As for presynaptic effects of GABABR activation in E*→*E synapses, the same study used a conditioning burst of 5 pulses at 50 Hz followed by a single test pulse to determine the time course of heterosynaptic short-term depression of excitatory postsynaptic currents [Isaacson et al., 1993]. This heterosynaptic short-term depression was shown to be GABABR-dependent, and it had a maximal amplitude (minimal ratio between the two stimuli) at approximately 300 ms and returned to baseline (ratio between the two stimuli close to one) at approximately 1600 ms [Isaacson et al., 1993]. Comparable PPR dynamics were measured for the GABABR-dependent short-term depression of inhibitory postsynaptic currents in CA1 pyramidal cells [Olpe et al., 1994]. In this work, we assigned the differences between the decay times in GABABR-dependent activity of GIRK or VGCC channels to the prevalence of RGS proteins by allowing the RGS protein concentration to vary between different domains (postsynaptic spine, excitatory presynaptic terminal and inhibitory presynaptic terminal), while we kept the reaction rates equal in all domains.

We applied a multiobjective optimization strategy to constrain our model. We varied 9 different model parameters according to the boundary conditions shown in Table 2B. We introduced objective functions to 1) minimize the difference between the experimentally measured GIRK channel current and the GIRK conductance predicted at 16 time points (80-700 ms after stimulus onset), and 2) minimize the summed difference between GABAB-dependent PPRs measured for excitatory and inhibitory presynaptic terminals and the PPRs predicted based on beta/gamma subunit of G_i_ binding to VGCCs — see Methods for details. We employed NSGA-II [Deb et al., 2002, Bahl et al., 2012] for 25 generations (population size N=2000) in 15 independent trials and for each trial picked the parameter sets that resulted in smallest normalized sum of pre- and postsynaptic objective functions.

The evolution of the objective functions showed little improvement after 10 generations (Fig. 1A–B). The best parameter set fit well into both post- (Fig. 1C) and presynaptic (Fig. 1D–E) data — see Table S1 for the chosen parameter set as well as the best parameter sets obtained from all 15 trials.

**Figure 1:**
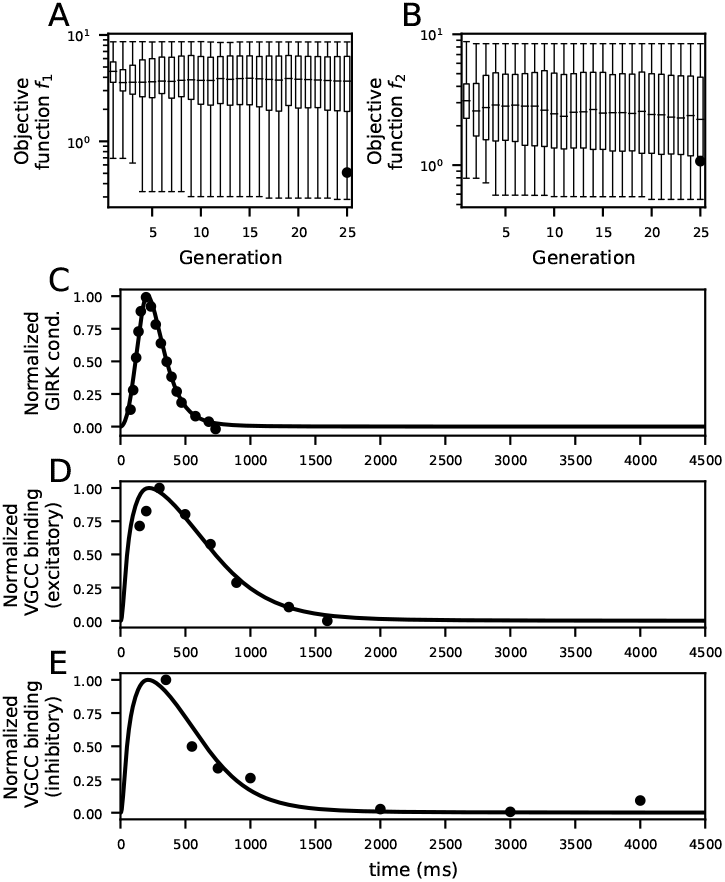
Fitting of the GABABR signalling pathway model. **A–B**: Evolution of the objective functions *f*_1_ (A) and *f*_2_ (B) across 25 generations of multi-objective optimisation. The parameter set chosen for simulations shown in panels (C)–(E) first appeared at the final (25th) generation (objective function values denoted by black dots) of the multi-objective optimisation — this parameter set had the smallest sum of objective functions normalized by their medians 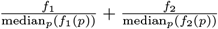. **C–E**: Time course of the experimental data-derived (dots) and model-predicted (solid lines) GIRK conductances at the postsynaptic neuron (C), VGCC binding at the presynaptic side of the E*→*E synapses (D), and VGCC binding at the presynaptic side of the I*→*E synapses (E).

Taken together, our model fits electrophysiological data on the GABABR-mediated short-term depression and postsynaptic GIRK conductance dynamics.

### 3.2 Verification of the presynaptic GABABR model by inclusion of additional components

While the postsynaptic GIRK conductance, as predicted by the model, can be directly related to measured electrophysiological data, the relationship between the predicted binding of the G_i*βγ*_ to the VGCCs and the PPR involves mechanisms not described in the model fitted in Fig. 1. Here, we aimed to make sure that the GABABR-activation-mediated binding of the G_i*βγ*_ to the VGCCs produced an effect that was rapid and strong enough to dampen the amplitude of the N-type Ca^2+^ currents in a realistic setting that involved the interplay of ionic currents in the presynaptic terminal.

We verified the appropriate functioning of the presynaptic model by inserting the biochemical model (Table 2) into a composite model of a piece of axon and a presynaptic terminal containing Hodgkin-Huxley-type description of Na^+^ and K^+^ currents (from [Hay et al., 2011]) as well as N-type Ca^2+^ currents (from [Migliore et al., 1995]), where the Ca^2+^ current kinetics were dependent on the binding of the G_i*βγ*_ to the N-type Ca^2+^ channel as reported in [Colecraft et al., 2000, Huynh et al., 2015] (Fig. 2A). We also incorporated a Ca^2+^-influx-mediated neurotransmitter release [Destexhe et al., 1994]. We assumed that the activation of the axon terminal either directly (in the I*→*E synapse) or indirectly (in the E*→*E synapse) caused GABA to be released and detected by the GABABRs in the axon terminal, as suggested by experimental data [Olpe et al., 1994, Isaacson et al., 1993, Scanziani, 2000]. To model the stimulation protocol of the heterosynaptic short-term depression of [Isaacson et al., 1993], the axon was stimulated only for the second incoming stimulus in the E*→*E synapse while the first stimulus (“S1” in [Isaacson et al., 1993]) was assumed to affect other nearby axon terminals that caused the GABA to be released and to activate the GABABRs in the modelled axon terminal. By contrast, to model the stimulation protocol of [Olpe et al., 1994] in the I*→*E synapse, the modelled axon terminal was activated for both stimuli, and since the GABABRs were assumed to function as autoreceptors, both stimuli triggered GABABR activation.

**Figure 2:**
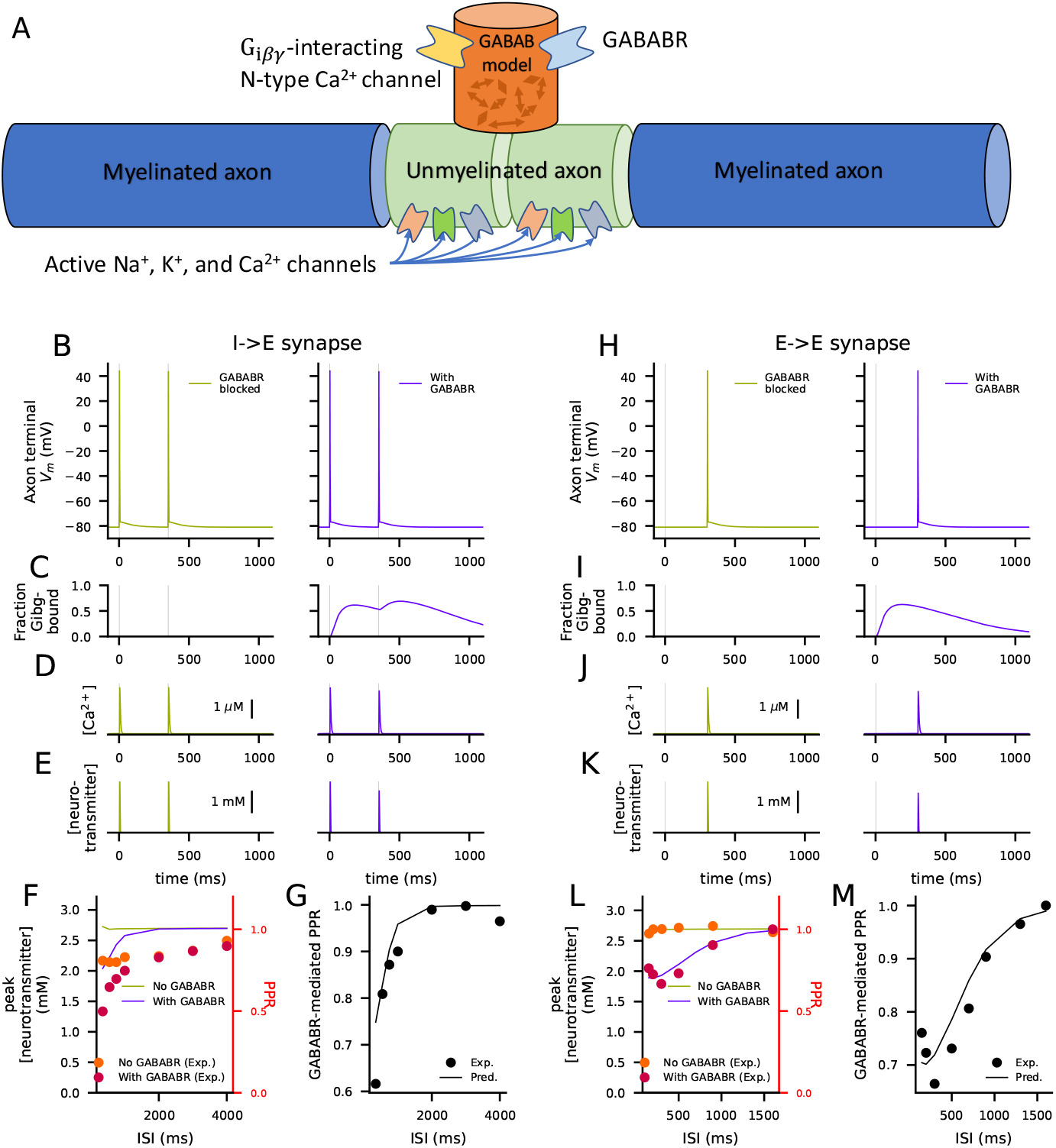
A model of a piece of axon with active ion channels for AP generation and the presynaptic terminal with the intracellular GABABR activation model coupled to N-type VGCCs reproduces the experimental PPR data. **A**: An illustration of the model of the presynaptic terminal. The model consists of two passive axonal compartments surrounding two active axonal compartments with Na^+^, and K^+^, and Ca^2+^ ion channel models from [Hay et al., 2011]. A presynaptic terminal with N-type VGCCs [Migliore et al., 1995] coupled to the intracellular GABABR activation model branched from between the active compartments. **B–D**: The membrane potential (B), the fraction of N-type VGCCs in the G_i*βγ*_-bound state (C), and the intracellular Ca^2+^ concentration (D) at the inhibitory presynaptic terminals with (blue) and without (light green) GABABR activation in response to two stimuli (at 0 and 350 ms) injected at the further end of the passive axon compartment. In the presence of GABABR-activation, the first stimulus activates the GABABRs, leading to inactivation of some of the N-type Ca^2+^ channels in the terminal (C, blue), which causes a depression of the Ca^2+^ transient induced by the second stimulus (D, blue). **E**: The neurotransmitter concentration at the synaptic cleft in response to the stimuli in the presence (blue) and absence (light green) of GABABR activation. The depression of the Ca^2+^ transient induced by the second stimulus leads to less neurotransmitter being released when GABABRs are present (E, blue). **F**: The peak of the neurotransmitter concentration (mM; right axis) corresponding to the second stimulus with respect to the inter-stimulus interval (ISI) between the two stimuli. The curves show the predicted peak amplitudes in the presence (blue) and absences (light green) of GABABR activation, and the data points indicate the PPRs (right axis) measured experimentally in the presence (orange) and absence (red) of GABABR antagonist. **G**: The peak of the neurotransmitter concentration of the GABABR-active case normalized by the corresponding peak in the GABABR-blocked case with respect to the ISI. The data points show the experimentally measured PPR in the GABABR-active case normalized by the corresponding PPR in the GABABR-blocked case. **H–K**: The experiment of (B–E) repeated for excitatory synapses with inter-stimulus interval 300 ms, but the first stimulus (“S1” at 0 s) only activates the GABABRs in the presynaptic terminal, not the AP generation in the axon. **L–M**: The experiment of (F–G) repeated for excitatory synapses.

In simulations of both I*→*E and E*→*E synapses, stimulation of the axon led to an AP in the axon terminal both in presence and absence of GABABR activation (Fig. 2B,H). The GABA inputs (indicated by gray vertical lines) caused activation of the presynaptic GABABRs, leading to G_i*βγ*_ binding to the presynaptic Ca^2+^ channels (Fig. 2C,I) and inactivating them. This decreased the amplitudes of the presynaptic Ca^2+^ transients (Fig. 2D,J) and the amplitude of the neurotransmitter release (Fig. 2E,K) in response to the second input (blue), while the neurotransmitter release was as strong for the second input as for the first input in the absence of GABABR activation (green). We repeated these simulations for all inter-stimulus intervals used in experimental setups [Olpe et al., 1994, Isaacson et al., 1993, Scanziani, 2000], and estimated the relative difference in the amplitude of neurotransmitter release between the first and the second electrical stimulation in each case. Although the absolute PPRs estimated as the relation of the neurotransmitter release between the second and first input was mildly overestimated both in presence and absence of GABABR activation in the I*→*E synapse (Fig. 2F), normalization of the GABABR-involving PPR curve by the GABABR-excluding one gave a good fit to the corresponding experimental data [Olpe et al., 1994] (Fig. 2G). The model of E*→*E synapse, by contrast, fit both absolute (Fig. 2L) and GABABR-normalized (Fig. 2M) heterosynaptic short-term depression curves measured in [Isaacson et al., 1993].

We ran a series of additional simulations of the composite model to show the validity of our model in describing the GABABR-mediated form of short-term synaptic depression. The results shown in Fig. 2 were produced with a representative parameter set out of the 15 tested ones, but the good fit was retained by most of the fitted parameter sets — in only 5/15 models the average error between predicted and measured normalized PPR curve was larger than 0.05 (Fig. S1A–B). We also analyzed the behaviour of our model under a stimulation protocol [Isaacson et al., 1993] that exhibited short-term facilitation of excitatory postsynaptic currents. Short-term facilitation index (i.e., the amplitude of the second EPSC response normalized that of the first one given two consecutive stimuli) is expected to be increased by processes or manipulations that decrease neurotransmitter release, such as GABABR-mediated inhibition of presynaptic VGCCs [Isaacson et al., 1993]. The validity of our biochemically detailed model in describing short-term plasticity was supported by these simulations: our model predicted that the facilitation index (here quantified as the ratio of neurotransmitter peak values) was 13.7% smaller in the absence compared to the presence of GABABR activation while experimental data suggests a decrease of 10.5*±*2.8% [Isaacson et al., 1993] (Fig. S1C–E).

Taken together, our model of GABABR-mediated presynaptic short-term depression fits the electrophysiological data also when included in a composite model including a biophysically detailed description of neurotransmitter release.

### 3.3 The model fitted to AP firing activity profiles from L5PCs

The data from [Isaacson et al., 1993] allowed fitting the parameters of the biochemically detailed model that govern the kinetics of the postsynaptic GIRK channel activation, but it remains to be shown that the model can correctly capture the effects of GABABR activation on input integration and AP generation in cortical cells. Here, we used a previous estimate of the single-channel conductance of the GIRK channel (33 pS [Chen and Johnston, 2005]) and data from [Schulz et al., 2021] where the AP firing profiles, i.e., the spiking response of L5PC to a combination of somatic and apical dendritic stimuli (5*×*6 data points) was recorded in presence and absence of GABABR activation. We then implemented our biochemically detailed model of GABABR pathway activation (Table 2) in a multicompartmental model of an L5PC [Hay et al., 2011] (illustrated in Fig. 3A) to compare the simulated effects of postsynaptic GABABR activation with the experimentally measured ones.

**Figure 3:**
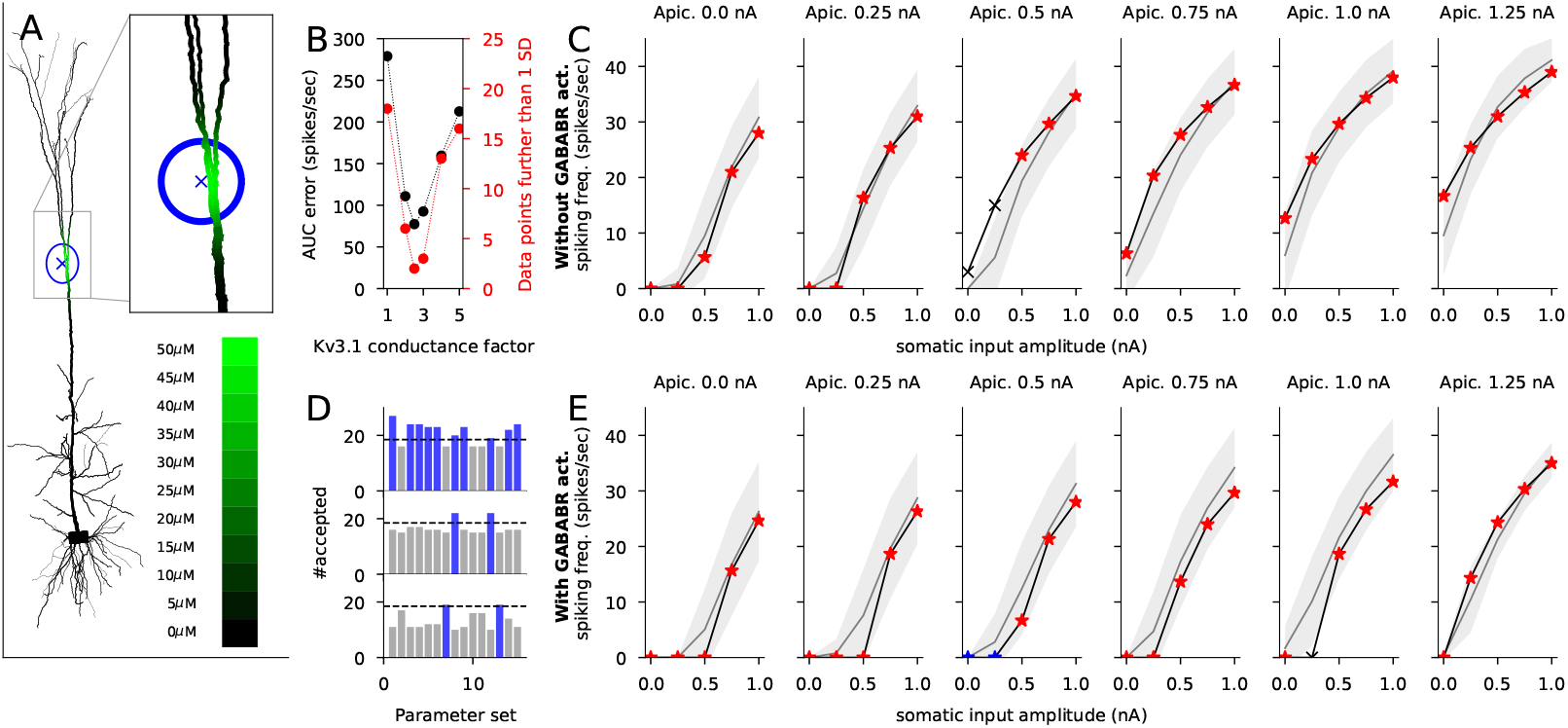
The model reproduces the GABABR-activation-dependent inhibition of the spiking response to apical and axo-somatic stimulation in L5PCs. **A**: The morphology of the L5PC model (from [Hay et al., 2011]; apical dendrite extending upwards from the soma, basal dendrites to other directions) and the GABABR agonist concentration at different compartments (color-coded). The locus of the simulated GABABR agonist puff was at the main branching point of the apical dendrite (650 µm up from the soma), the maximal agonist concentration was 50 µM, and the SD of the agonist concentration was 45 µm. **B**: The fitting of Kv3.1 conductance. The black data (left-hand side y-axis) shows the summed error of firing rate in response to a combination of axo-somatic (0.0, 0.25, 0.5, 0.75, or 1.0 nA; see (C)) and apical dendritic (0.0, 0.25, 0.5, 0.75, 1.0, or 1.25 nA; see (C)) DC stimulus between the predicted and experimentally measured [Schulz et al., 2021] data given different scaling factors (x-axis) of the Kv3.1 conductance in the absence of GABABR activation. The red data (right-hand side y-axis) show the number of predicted data points that were within 1 SD from the measured data. A scaling factor of 2.5 gave the smallest difference between model prediction and experimental data both in terms of total error (black) and number of correctly predicted data points (red) and was thus used henceforth. **C**: The predicted (black curve with markers) and experimentally measured (gray curve with shaded area showing the mean *±* SD) firing rates in response to the combined axo-somatic and apical dendritic stimulation in the absence of GABABR activation. The x-axes show the amplitude of the axo-somatic DC stimulus, while the amplitude of the apical dendritic DC stimulus (0.0–1.25 nA) is indicated above the plots. The red asterisks and the black crosses indicate the model predictions that were closer or further, respectively, than 1 SD from the experimentally measured firing rate [Schulz et al., 2021]. The model correctly predicted the firing rate for 28/30 stimulus combinations. **D**: The number of correctly (prediction within 1 SD from experimental data) predicted firing rates in the presence of GABABR activation using reaction-rate parameter sets from 15 different GABABR activation model fits (see Table S1). The blue bars label the parameter sets that correctly predicted the firing rate in the presence of GABABR activation for two thirds (19/28) or more of the stimulus combinations that yielded acceptable predictions in the absence of GABABR activation in (C). By contrast, the gray bars indicate parameter sets that correctly predicted the firing rate in the presence of GABABR activation for 18 or fewer stimulus combinations. The upper panel shows the results when a 45 µm SD of GABABR agonist distribution was used, yielding optimal fit to the data, while the middle and lower panels show the results where a too small (20 µm) or too large (100 µm) SD was used. **E**: The predicted (black curve with markers) and experimentally measured (gray) firing rate in response to the combined stimulus in the presence of GABABR activation; see (C) for the GABABR-blocked case. The model correctly (within 1 SD) predicted the firing rate for 27 out of the 28 stimulus combinations for which the GABABR-blocked model correctly predicted the firing response, see (C). The model also correctly predicted the firing rate for two stimulus combinations for which the GABABR-blocked model failed to predict the correct firing response (blue asterisks).

We first noticed that the Hay model did not fit well with the experimental data collected in the absence of GABABR activation [Schulz et al., 2021] — instead, the predicted firing frequencies were constantly lower than observed in the data. Preliminary analyses where the conductance of a single type of ion channels was altered showed that increase of the Kv3.1 channel conductance, but not other alterations of ion-channel conductances, changed the AP firing profile of the model L5PC toward the data measured from (non-GABABR-activated) L5PCs in [Schulz et al., 2021]. Although the Kv3.1 channels are hyperpolarizing, they exhibit a rapid channel deactivation that promotes fast repetitive firing in most neuron types, and they are expressed in L5PCs [Ichinohe et al., 2004, Kaczmarek and Zhang, 2017]. The best fit to the data from [Schulz et al., 2021] was obtained by using the Hay model where the Kv3.1 channel conductance was 150% larger than in the standard model: with this factor 28/30 of the predicted firing frequencies were within 1 standard deviation (SD) from those measured in [Schulz et al., 2021] (Fig. 3B–C).

We next implemented the biochemically detailed model of GABABR activation (Table 2) and its coupling to a GIRK channel model, adapted from [Yim et al., 2015], in the L5PC model. We repeated the simulations of combined somatic and apical dendritic stimuli in the presence of GABABR activation using the 15 optimized biochemical models. In [Schulz et al., 2021], it was estimated that the GABABR agonist (50 µM) was effective up to a distance of 100 µm from the injection point at the apical dendrite 500–800 µm from the soma. To model this, we used a Gaussian density function centered at 650 µm toward the apical dendrite from the soma — the GABABR agonist concentration was set 50 µM at the centre and decreased with an SD of 20, 45, or 100 µm (Fig. 3A). It should be noted that we did not take into account the possible contribution of dendritic VGCCs to this inhibition — see Discussion. We found that the simulations performed with GABABR agonist distribution with a SD of 45 µm fit well to the spiking data from [Schulz et al., 2021]: 29 of 30 somatic-apical stimulus combinations (including 27 of the 28 that were correctly reproduced in the absence of GABABR activation) were correctly reproduced, i.e., such that the predicted value was within 1 SD from the value measured in [Schulz et al., 2021], in the presence of GABABR activation (Fig. 3D–E). This was a relatively robust result, since for 10/15 optimized parameter sets, two thirds (19/28) or more of the somatic-apical stimulus combinations were correctly reproduced in the presence of GABABR activation.

Taken together, our model fits postsynaptic spiking data from L5PCs measured in the presence of GABABR agonist well. Next, we use our model to explore the pre- and postsynaptic effects of GABABR activation in L5PCs where activation is entirely driven by synaptic inputs.

### 3.4 The effects of pre- and postsynaptic GABABR activation on L5PC firing induced by in vivo-like synaptic inputs

Our common framework for modelling pre- and postsynaptic GABABR activation offers a unique opportunity to compare the effects of the two. Here, we modeled the effects of both pre- and postsynaptic GABABR stimulation on dendritic membrane potential and spiking activity during glutamatergically driven activity in an L5PC. To do this, we fit our model to in vivo patch-clamp-recorded data from L5PCs during electrical paw stimulation [Palmer et al., 2012]. In [Palmer et al., 2012], short electric pulses administered at the rat paw induced a dense spiking activity in L5PCs for 0.6 seconds. This activity was significantly weakened by GABABR activation mediated by interhemispheric inputs [Palmer et al., 2012]. We fit the model parameters pertaining to L5PC activity induced by this stimulation protocol (number of excitatory synapses and GABA flux at dendritic compartments) and used the obtained model for exploring the contributions of pre- and postsynaptic GABABR activity to L5PC firing. Note that we here did not model the effects of GABABR activation on postsynaptic voltage-gated Ca^2+^ channels although they, according to the results of [Palmer et al., 2012], mediate part of the GABABR-driven interhemispheric inhibition of L5PCs.

The L5PC postsynaptic response evoked by paw activation in [Palmer et al., 2012] had a dynamic structure, where peaks and troughs of inputs alternated approximately every 50 ms for a total time of 600 ms. To model this, we stimulated 20 independent axons, modelled as in Section 2.2, every 50 ms twelve times with a probability of 0.5 (i.e., each axon fired on average six times). A number (*N*_syn_) of synapses using the neurotransmitter-gated AMPA/NMDA model (Section 2.5) were distributed across the distal half of the apical dendritic tree of the L5PC (distance from soma *>* 650 µm, longest branch length being 1300 µm), and each was coupled to the neurotransmitter release model corresponding to one of the 20 (randomly chosen) presynaptic axon terminals. We repeated this stimulation 10 times with an interval of 10 seconds which was adequate for the L5PC to return to baseline. We added a stimulating somatic current lasting throughout the 600-ms periods whose amplitude was randomly picked (independently for each trial) from a uniform distribution between 0.15 and 0.25 nA — this produced a trial-to-trial variability of neuronal spike output, similar to [Palmer et al., 2012]. To model the interhemispheric inhibition of [Palmer et al., 2012], we inserted GABABRs and GIRK channels activated by them to the distal half of the apical dendritic tree.

We first adjusted the amount of excitation in a single L5PC (number of glutamatergic synapses, Fig. 4A) in the absence of GABABR activation (see Fig. 4B) to fit to the firing rate (2.9 spikes/sec) during the high-activity period. The best fit was obtained using 500 glutamatergic synapses (Fig. 4C) — i.e., each axon was connected to 25 synapses randomly placed on the distal apical dendrite. We then fit the amount of GABA flux to the GABABRs to reproduce the lower firing frequency (2.2 spikes/sec) in the presence of GABABR activation. Following the observations of [Palmer et al., 2012], we first excluded the GABABR activation from the presynaptic terminals. The data from [Palmer et al., 2012] was reproduced with a GABA flux of 3.25 nM/ms (Fig. 4D). The spiking patterns in the presence and absence of postsynaptic GABABR activation are illustrated in Fig. 4E–F.

**Figure 4:**
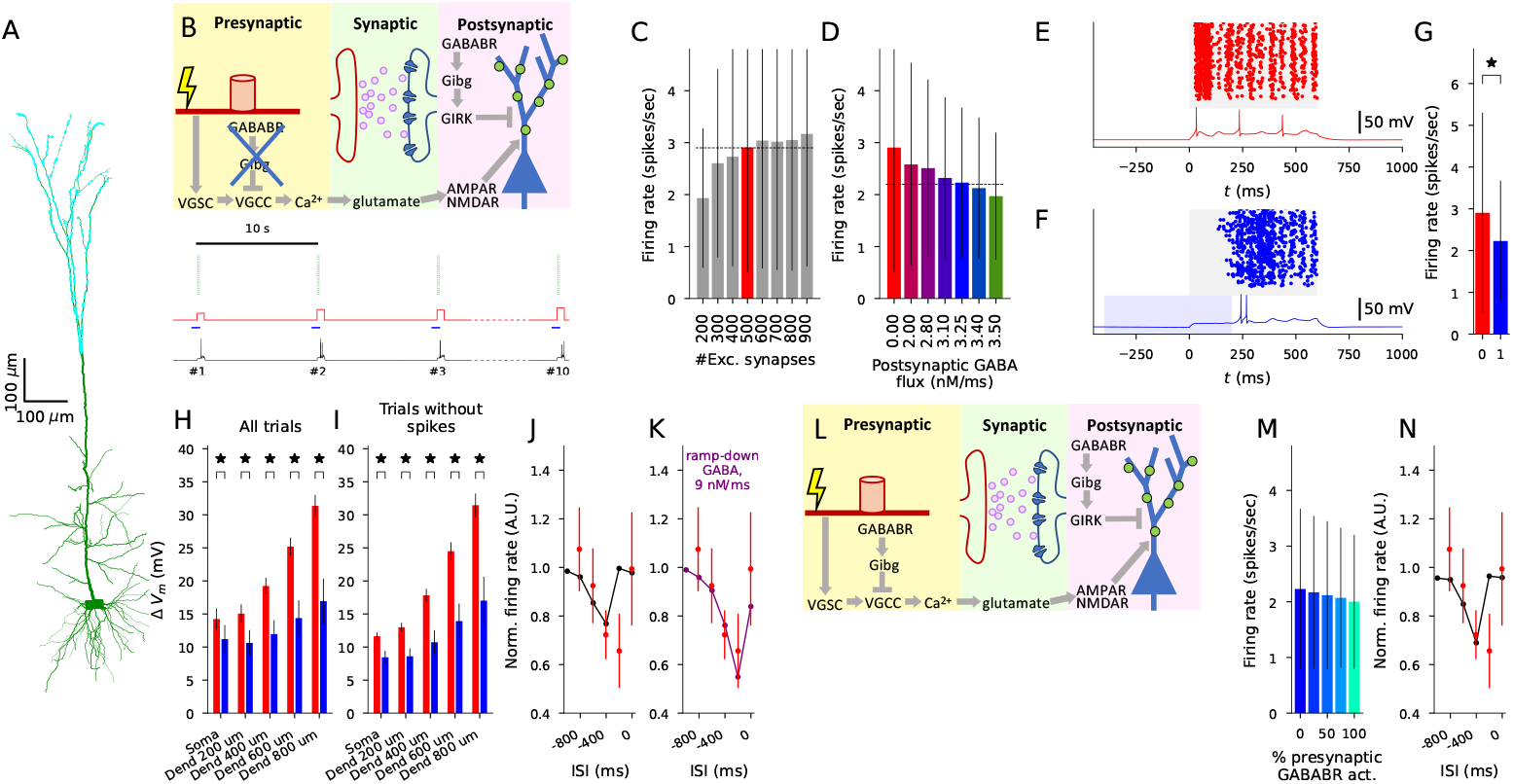
The model can be fitted to in vivo data on interhemispheric inhibition to explore the GABABR-mediated suppression of high firing activity in L5PCs. **A**: Illustration of the dendritic tree of the modelled L5PC [Hay et al., 2011] (green) and the glutamatergic inputs (N=500) distributed across the distal half of the apical dendrite (cyan). **B**: Illustration of the GABABR actions and the stimulation. Top: Presynaptic stimulation leads to activation of axonal voltagegated Na^+^ channels (VGSCs), which opens the VGCCs, leading to glutamate release. The glutamate activates postsynaptic AMPARs and NMDARs, leading to depolarization of the membrane. When postsynaptic GABABRs are activated, the GIRK channels counteract this by hyperpolarizing the membrane. In panels (C)–(I), the presynaptic GABABRs are inactive and all VGCCs are in the “willing” state. Bottom: Illustration of the stimulation of the 20 axons (green), the concurrent subthreshold stimulation of postsynaptic L5PC soma (red), the periods of GABABR activation (blue), and the resulting somatic membrane potential trace (black). **C**: The firing rate (spikes/sec) following the stimulus onset in the absence of postsynaptic GABABR-activating inputs given different numbers of glutamatergic synaptic inputs. The stimulus consisted of 600 ms of rhythmic, 20 Hz stimulation of the axons where the probability of each stimulus was 0.5. The high-frequency stimulation was combined with a synchronous and equally long (600 ms) DC at the soma with an amplitude randomly picked from a uniform distribution between 0.15 and 0.25 nA. We simulated L5PCs with 40 different randomly picked distributions of synapses, and each L5PC was given 10 independent stimuli administered every 10 seconds (i.e., each simulation lasted 100 seconds, plus a 10.6 second period for reaching a steady state and recording the output), leading to 400 trials. The data of [Palmer et al., 2012] was best reproduced by using N=500 synapses. **D**: The firing rate (spikes/sec) following the stimulus onset in the presence of postsynaptic GABABR-activating inputs given different fluxes of GABA. The GABABR model (Table 2) was included in each compartment in the distal half of the L5PC. The onset of the GABA input was 400 ms before that of the glutamatergic inputs and lasted for 600 ms. The data of [Palmer et al., 2012] was best reproduced by using the GABA flux of 3.25 nM/ms. **E–F**: Illustration of the L5PC spiking across all trials and membrane potential time course of a single trial during the high-activity period (starting at *t* = 0 ms) in the absence (E) and presence (F) of the GABA flux of 3.25 nM/ms. **G**: The L5PC firing rate during the high-activity period was significantly lower in the presence (blue) than in the absence of the postsynaptic GABABR-activating inputs (U-test, p=0.022*<*0.05). **H–I**: The average membrane potential during the high-activity period in the absence (red) and presence (blue) of the postsynaptic GABABR-activating inputs. The asterisks denote statistical significance between absence and presence of postsynaptic GABABR-activating inputs (p*<*0.005) when averaged across all trials (H) or the trials where no APs were induced (I). **J**: The normalized firing rate following the stimulus onset in the presence of postsynaptic GABABR-activating inputs with varied time of onset. The firing rates were normalized by the corresponding firing rate in the absence of GABABR-activating inputs. The red dots show the data measured in [Palmer et al., 2012], while the black curves show the model predictions. **K**: The normalized firing rate following the stimulus onset in the presence of a ramp-down form of postsynaptic GABABR-activating inputs with varied time of onset. See Fig. S2 for details. **L**: Illustration of the stimulation and the GABABR actions. In panels (M)–(N), the presynaptic GABABRs were activated in all or some of the 20 axons in addition to the postsynaptic GABABR actions. **M**: The firing rate following the stimulus onset in the presence of pre- and postsynaptic GABABR-activating inputs. In these simulations, the presynaptic terminal was given the same GABABR-activating inputs as the postsynaptic GABABR-containing dendritic compartments, leading to the inactivation of some of the presynaptic N-type Ca^2+^ channels as in Fig. 2. The fraction of presynaptic terminals (randomly picked among the 20) to which GABABR-activating ligands were added is varied from 0% (the data from blue bars of panels (D) and (G)) to 100% (all presynaptic terminals were given GABA inputs). **N**: The experiments of panel (J) repeated by administering both pre- and postsynaptic GABABR-activating inputs as in (M).

Next, we tested whether the difference in the numbers of APs and dendritic membrane potentials between the absence and presence of GABABR activation was significant. Similar to [Palmer et al., 2012], the firing in the presence of GABABR activation was significantly decreased compared to the absence (U-test, p=0.02*<*0.05; N = 400, 400; Fig. 4G). However, contrary to [Palmer et al., 2012], the average membrane potential during the 600 ms following the simulated paw stimulation was also significantly decreased in the presence of GABABR activation compared to the absence (Fig. 4H–I). This was the case both when considering all trials (Fig. 4H) and when considering only the non-spiking trials (Fig. 4I), and the result held both for somatic membrane potentials as well as membrane potentials recorded at four different locations along the apical dendrite (U-test, p*<*0.005, i.e., 0.05 Bonferroni-corrected by multiple (10) testings). We also tested the effect of the timing of the GABABR activation on the L5PC firing during the high-activity period. As observed in [Palmer et al., 2012], our model predicts that GABABR activity that is either synchronous with the glutamatergic activity or preceding it by too much (GABABR-activity onset *≥* 800 ms before the onset of the glutamatergic inputs) does not suppress the L5PC firing during the high-activity period (Fig. 4J). For an interval of 200 ms, however, our model predicted normal firing rate while the data of [Palmer et al., 2012] suggested a strong suppression of L5PC firing during the high-activity period. By contrast, a strong suppression for the 200 ms interval was obtained if we modified the simulation to use a ramp-down GABA stimulus instead of the square-pulse GABA stimulus of Fig. 4D–J (Fig. 4K; Fig. S2).

We next explored the role of presynaptic GABABR activation in inhibiting L5PC activity (Fig. 4L). First, we carried out simulations where the presynaptic VGCCs were externally set to be maximally G_i*βγ*_-bound regardless of the GABA inputs, and the postsynaptic GABABR activation was the same as before (a square-pulse flux of 3.25 nM/ms for 600 ms). If we assumed all VGCCs to be G_i*βγ*_-bound, we achieved a firing rate of 1.58 spikes/sec (difference to postsynaptic-only GABABR activation was Δ*f* = -0.65 spikes/sec) during the high-activity period, whereas if we chose a quasi-maximal level of G_i*βγ*_-bound VGCCs, that is, a steady state implied by the assumption of all G_i_ being dissociated and the ratio of the forward and backward rates of G_i*βγ*_ binding to VGCC (which resulted in 85.7% of VGCCs being G_i*βγ*_-bound), we achieved a firing rate of 1.75 spikes/sec (Δ*f* = -0.48 spikes/sec; Fig. S3A). The effects of the presynaptic VGCC inhibition were not linearly cumulative with those of the postsynaptic GABABR activation: when blocking the postsynaptic GABABR activity, the inclusion of VGCC inhibition decreased the firing rate from 2.9 spikes/sec only to 2.66 spikes/sec (Δ*f* _no postsyn. GABABR_ = -0.24 spikes/sec) or 2.71 spikes/sec (Δ*f* _no postsyn. GABABR_ = -0.19 spikes/sec) in the maximal and quasi-maximal modes of VGCC inhibition, respectively (Fig. S3A).

Finally, we tested whether similarly large effects were obtained by applying presynaptic GABA inputs similar to those affecting the postsynaptic GABABRs (a square-pulse flux of 3.25 nM/ms for 600 ms). Depending on the fraction of presynaptic terminals affected by GABA, the firing rate of the L5PC was 2.0–2.17 spikes/sec (Δ*f* = -0.06 – -0.23 spikes/sec; Fig. 4M). These changes were statistically non-significant, concordant with the observations of [Palmer et al., 2012]. The presence of presynaptic GABABR activation also had little effect on the sensitivity of the L5PC firing to the timing of the GABABR-activating stimulus (Fig. 4N). We also tested if presynaptic GABABR activation could have larger effects on the L5PC firing if its internal mechanisms and the GABA binding were enhanced. To do this, we increased the GABA flux and decreased the RGS concentration in the presynaptic terminals. However, these changes had mild strengthening effects of the GABABR-mediated inhibition of the L5PC firing: either increasing the presynaptic GABA flux 2-to 4-fold (Fig. S3B) or decreasing the presynaptic RGS concentration by 25–90% (Fig. S3C) only decreased the firing during the high-activity period to 1.96 – 2.0 spikes/sec (Δ*f* = 0.23 – 0.27 spikes/sec).

Taken together, the predictions obtained with our model fitted to experimentally measured features of interhemispheric inhibition highlight the role of postsynaptic rather than presynaptic GABABR activation and suggest that the GABA input activating the postsynaptic GABABRs is strong shortly after the onset of the ipsilateral stimulation and weaker afterwards. While our model predicts that by completely inhibiting presynaptic VGCCs can significantly add to the interhemispheric inhibition mediated by postsynaptic GABABR activation, it also suggests that such a degree of VGCC inhibition cannot be achieved by presynaptic GABABR activation alone.

### 3.5 Model-aided analysis of RGS effects on neuronal dynamics

The biochemically detailed nature of our model permits using it for explaining and studying the implications of data from genetic knockout experiments. In addition to the proteins that directly mediate the effects of GABABR activation, such as the GIRK channels, VGCCs, and the GABABRs themselves, an important class of proteins included in our model are the RGS proteins. A primary role of RGS proteins, especially the R7 family (RGS6, RGS7, RGS9, and RGS11), is to terminate the G-protein activity by facilitating the GTPase activity of the G*α* subunit, but they also mediate other G-protein-related actions through forming a complex with other proteins [Anderson et al., 2009]. In a genetic knockout experiment [Ostrovskaya et al., 2014], the authors showed that RGS7 contributed to termination of GABABR-mediated postsynaptic GIRK currents in CA1 pyramidal neurons, but they also reported an RGS7-mediated increase in the sensitivity of GIRK channels to GABABR activation. Namely, RGS7-KO neurons responded to approximately 5 times smaller GABABR agonist concentrations than wild-type neurons [Ostrovskaya et al., 2014]. Importantly, the authors also reported that the RGS7-KO altered the CA1 neuron excitability and the amplitude of low-frequency stimulation (LFS) induced LTD [Ostrovskaya et al., 2014]. The observed effects could have been a direct consequence of the decreased amount of RGS proteins, but they could also stem from increased sensitivity of GIRK channels to GABABR activation. While the prolonged termination of GIRK currents is likely to be a direct consequence of the former, it remains unknown whether changes in firing threshold and plasticity are created by the former or the latter cause. Here, we showcase the potential of our model in answering mechanistic questions such as this one by analysing the effects of different manipulations of the GABABR system in a multicompartmental CA1 pyramidal neuron model [Combe et al., 2018].

We tested the effects of partial removal of RGS and the altered baclofen (GABABR agonist) sensitivity on the GIRK current deactivation in CA1 pyramidal cells (Fig. 5A, green). This was done in the absence of synaptic stimulation. A 90% reduction in RGS concentration caused the same magnitude of prolongation of the GIRK current activity (≈10 s) as observed in RGS7-KO [Ostrovskaya et al., 2014] (Fig. 5B), suggesting a minor contribution of RGS protein other than RGS7 to GIRK current termination in CA1 pyramidal cells. However, 90% RGS reduction alone decreased the somatic resting membrane only by 0.01–0.6 mV depending on the basal level of GABABR activity (Fig. 5C). By contrast, a 5-fold increase in GABABR agonist sensitivity (simulated by using 5-fold larger concentration) decreased the resting membrane by 1.0–3.2 mV or 1.3–3.8 mV (depending on the basal level of GABABR activity) when unaccompanied or accompanied, respectively, by the 90% RGS reduction (Fig. 5D–E). Likewise, the 90% RGS reduction alone had little effect on the firing threshold in response to somatic stimulus (Fig. 5G), but a 5-fold increase in GABABR agonist sensitivity, with (Fig. 5H) or without (Fig. 5I) the reduction in RGS concentration, increased the firing threshold by 14–18% or 15–51%, respectively, depending on the basal level of GABABR activity. The experimentally measured effects of RGS7-KO on resting membrane potential (−3.8 mV *±* 0.7 mV; Fig. 5F) and threshold current (140% *±* 12%; Fig. 5J) in CA1 pyramidal neurons were best approximated by our model with the 90% RGS reduction and a 5-fold increase in GABABR agonist sensitivity given a basal GABABR activity of approximately 10 A.U. (Fig. 5F,J; cyan).

**Figure 5:**
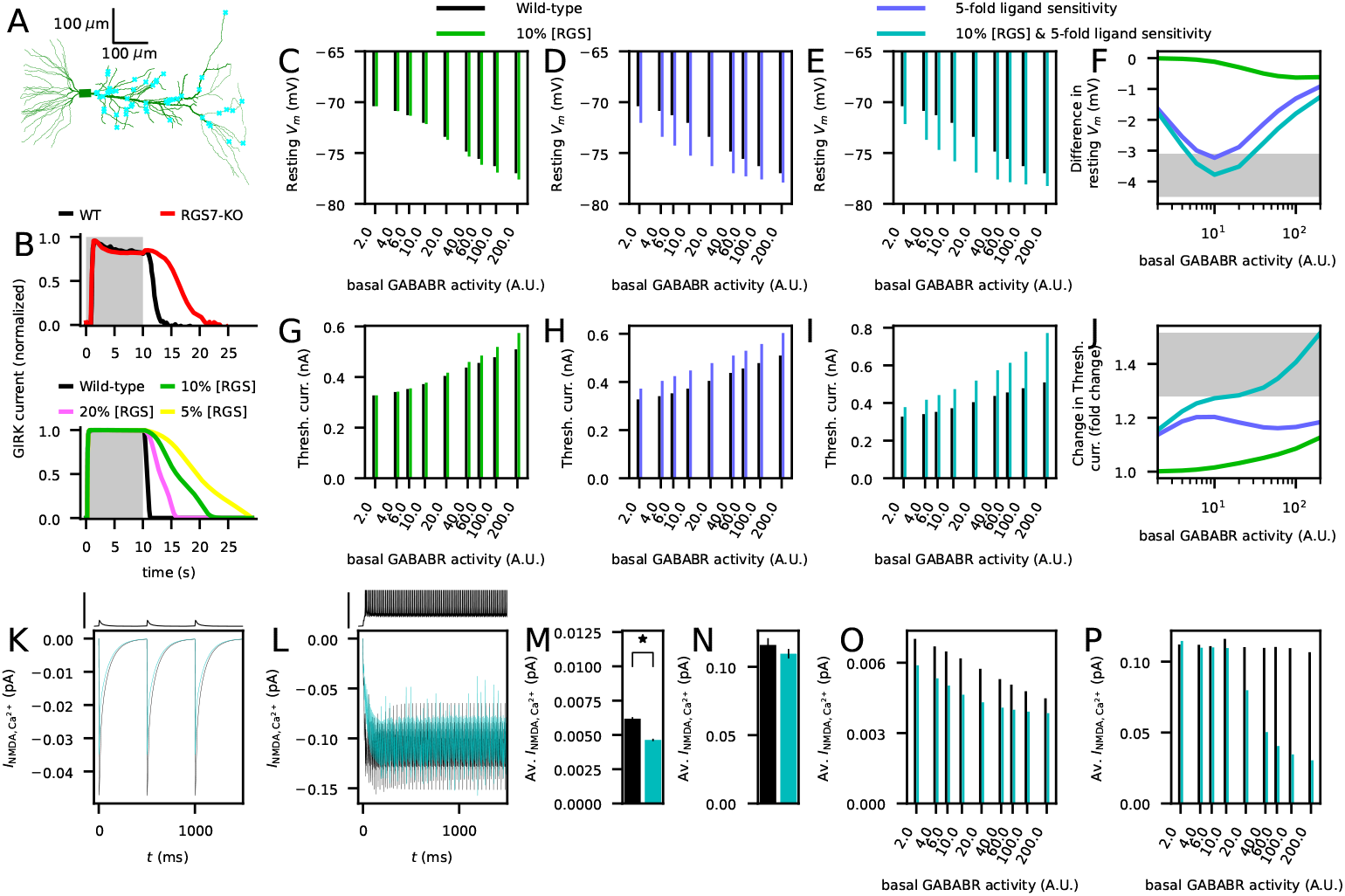
**Reduction of RGS concentration to 10% and 5-fold sensitivity to GABABR agonists prolongs GABABR-mediated GIRK currents, hyperpolarizes the membrane, increases the firing threshold, and decreases the Ca**^2+^ **transients induced by LFS in CA1 pyramidal cells in accordance with experimental data from an RGS7-KO**. **A**: The morphology of the Combe CA1 neuron model [Combe et al., 2018] (green) and an illustration of the 50 synapses randomly placed at the apical dendrite (cyan). **B**: Upper panel: Time course of baclofen-activated GIRK currents in CA1 pyramidal cells of wild-type (black) and RGS7-KO (red) mice (data extracted from [Ostrovskaya et al., 2014]). Lower panel: Model predictions for GIRK currents recorded at the apical dendritic compartment closest to soma in response to GABABR activation by a saturating agonist concentration (100 µM) at the default (Table 2B) RGS concentration (black) and when the RGS concentration was reduced to 20% (pink), 10% (green), or 5% (yellow). The RGS reduction by 90% (green) fit well the experimental observations (upper panel). The time of baclofen administration is indicated by the shaded areas. **C–E**: Model predictions for somatic resting membrane potential in the Combe model integrated with the GABABR model (i.e., the “wild-type” model) as a function of basal GABABR activity (modelled using a constant, non-depleting GABABR agonist concentration such that 1 A.U. corresponds to 1 pM agonist concentration). The black bars represent the Combe model integrated with the default GABABR model (Table 2), whereas the colored bars represent the candidate models for RGS7-KO; namely, the GABABR model where the RGS concentration was reduced by 90% (green; C), where the sensitivity to GABABR agonist was 5-fold larger (blue; D), or where both manipulations were present (cyan; E). **F**: The difference between the resting membrane potential in the wild-type model and the candidate models for RGS7-KO as a function of basal GABABR activity. The shaded area represents the mean *±* SD of the experimentally observed difference in resting membrane potentials between wild-type and RGS7-KO mice [Ostrovskaya et al., 2014]. **G–I**: Model predictions for the threshold current (for a somatic square-pulse current of 5 ms) in the wild-type model (black) and the candidate models of RGS7-KO (green, blue, cyan — see (C)–(E)). **J**: The ratio of threshold current in the wild-type model to that in the candidate models of RGS7-KO as a function of basal GABABR activity. The shaded area represents the mean *±* SD of the experimentally observed ratio of threshold currents between wild-type and RGS7-KO mice [Ostrovskaya et al., 2014]. **K–L**: NMDAR-mediated Ca^2+^ currents (averaged over the 50 synapses distributed across the apical dendrite) in response to the 2 Hz (K) or 100 Hz (L) stimulation. Black curves represent the response in the wild-type model and the cyan curves represent the candidate RGS7-KO model where RGS concentration was reduced by 90% and the sensitivity to GABABR agonist was 5-fold larger (averaged across N=5 simulations). The basal GABABR activity was 10 A.U. Insets above the graphs: Somatic membrane potential time course. Scale bar 50 mV. **M–N**: The average NMDAR-mediated Ca^2+^ currents in the 2 Hz (M) and 100 Hz (N) protocol. The NMDAR-mediated Ca^2+^ currents were first averaged across the stimulus interval (500 or 10 ms), and the mean across the 10 last stimuli were calculated. The bars show the mean and SD of N=5 simulations. In the 2 Hz protocol, the Ca^2+^ current amplitudes were significantly smaller (U-test, p=0.009*<*0.05) in the RGS7-KO model compared to the wild-type model in response to 2 Hz stimulus, but the difference in the Ca^2+^ current amplitudes in response to the 100 Hz stimulus did not reach significance (U-test, p=0.076*>*0.05). **O–P**: The mean NMDAR-mediated Ca^2+^ current amplitudes in the 2 Hz (O) and 100 Hz (P) protocol for the wild-type and RGS7-KO model across different basal GABABR activity levels. The RGS7-KO model predicts a difference in the HFS protocol only for large basal GABABR activity, whereas for the LFS protocol the Ca^2+^ current amplitudes are lower in the RGS7-KO model throughout the tested GABABR activity levels.

We next simulated the effect of RGS7-KO on Ca^2+^ inputs during stimulation protocols that induced plasticity in CA3-CA1 synapses. To do this, we distributed 50 AMPAR/NMDAR synapses [Wang, 1999, Mäki-Marttunen and Mäki-Marttunen, 2022] randomly across the apical dendrite of the CA1 pyramidal neuron model (Fig. 5A, cyan) and stimulated them rhythmically either at a 2 Hz (low-frequency stimulation; LFS) or 100 Hz (high-frequency stimulation; HFS). The NMDAR-mediated Ca^2+^ currents were significantly larger in HFS stimulation compared to LFS stimulation (Fig. 5K–P), owing to the induction of spiking in response to the HFS but not LFS protocol according to our model (Fig. 5K–L insets). Our model of RGS7-KO (90% RGS reduction in RGS concentration and 5-fold increase in GABABR agonist sensitivity) in moderate basal GABABR activity (10 A.U.) expressed significantly smaller NMDAR-mediated Ca^2+^ currents in response to LFS (Fig. 5K,M) but unaltered NMDAR-mediated Ca^2+^ currents in response to HFS (Fig. 5L,N), which is in accordance with the observations of decreased amplitude of LFS-induced long-term depression (LTD) and unaltered HFS-induced long-term potentiation (LTP) [Ostrovskaya et al., 2014]. However, our model suggested that in higher levels of basal GABABR activity the Ca^2+^ currents are decreased in response to both LFS (Fig. 5O) and HFS (Fig. 5P) in RGS7-KO.

Taken together, our model suggests that both a 90% reduction in RGS concentration and a 5-fold increase in GABABR agonist sensitivity are required for the effects of RGS7-KO on the GIRK current deactivation time, resting membrane potential and firing threshold in CA1 pyramidal cells. Our model also suggests that the difference between experimentally observed effects of RGS7-KO on LFS-induced LTD compared to HFS-induced LTP is due to the RGS7-KO significantly decreasing the NMDAR-mediated Ca^2+^ current amplitudes in response to LFS but not those in response to HFS.

## 4 Discussion

Here, we developed a biochemically detailed model of GABABRs and their effects on neuronal excitability through their G_i*βγ*_-mediated interactions with GIRK channels and VGCCs. Our model has two important advances compared to existing models. Firstly, the model describes the GABABR and G_i*βγ*_ activity in both pre- and postsynaptic terminals. The reaction rates were the same in different domains, only the concentrations of proteins (here, the RGS proteins) varied depending on the cell type and location. Thanks to this design principle, the model can be flexibly used in other cell types as well — only the concentrations of the proteins involved, especially the RGS proteins that regulate the duration of the GABABR-mediated effects, may need to be adjusted to brain-area-specific data (see [Mäki-Marttunen et al., 2020] for an example of such a task). Secondly, we described the mass-action-law-based interactions between GABABRs, G_i_ protein subunits, RGS proteins, and the target proteins (GIRK channels and VGCCs) instead of relying on simplified dynamical interdependencies. Although including these interactions makes the model somewhat heavier to simulate, it permits integration with larger models (e.g., models describing the effects of G_i*α*_ activation). Importantly, it also allows testing hypotheses on the effects of genetic manipulations of the involved proteins and, although not tested here, pharmacological manipulations. In comparison, the earliest models of GABABR-mediated K^+^ currents, like many newer models, treat these currents essentially the same way as GABAAR-conducted currents: a presynaptic AP induces a postsynaptic conductance with a predefined time course [Traub et al., 1994, Turi et al., 2019]. Of the previous biochemically detailed models, the model of [Destexhe and Sejnowski, 1995] describes the dynamics of the postsynaptic GABABRs and G-proteins (and how they activate K^+^ currents) in a simplified manner (no separate *α* or *β*-*γ* subunits and no binding to GIRK channels). In [Li et al., 2020], GABA activates presynaptic GABABRs and mediates short-term depression, but no G-protein interactions are described. To our knowledge, our model is the first to incorporate detailed data on the dependence of GIRK channel conductance [Dascal and Kahanovitch, 2015] and VGCC voltage-dependence and kinetics [Huynh et al., 2015] on G_i*βγ*_ binding into the GABABR model. This extends the biological realism of computational models of GABAergic inhibition and can thus lead to improved quantification of neuronal activity at multiple scales.

We applied our model to data from RGS7-KO [Ostrovskaya et al., 2014] to show the potential of our model in complementing experimental data to explain the mechanisms of genetic manipulations. In [Ostrovskaya et al., 2014], two key effects of RGS7-KO were reported: the decay time of GIRK conductance after a period of saturating level of the GABABR agonist baclofen was significantly extended, and the sensitivity to baclofen (in the domain of non-saturating concentrations) was increased 5-fold. While the first effect is likely to be a direct effect of decreased concentration of RGS proteins, the latter effect may be a subtype-specific interaction mediated by a heterotrimer composed of the *β*_5_ subunit of the G-protein, RGS7 (or RGS6, RGS9 or RGS11), and the RGS7-binding protein R7BP [Ostrovskaya et al., 2014] or another, yet unknown mechanism. The exact mechanisms underlying the latter effect are unclear — here, we applied a model where we used 5 times larger GABABR agonist concentration to effectively capture the increased baclofen-sensitivity of RGS7-KO reported in [Ostrovskaya et al., 2014]. We showed that both of these subcellular-level effects (decreased concentration of the G_i*α*_-GTP-hydrolysing RGS proteins and increased sensitivity to GABABR ligands) were needed to explain the electrophysiological phenotypes of the RGS7-KO, namely, the decreased resting membrane potential and the increased firing threshold. We also found out that the effects of RGS7-KO on plasticity in CA1 pyramidal cells (decreased LTD amplitude but unaffected LTP amplitude [Ostrovskaya et al., 2014]) are likely to be a direct consequence of the stimulus-induced postsynaptic Ca^2+^ flux being significantly reduced in the RGS7-KO model during LFS but not HFS. Furthermore, we showed that the prolongation of GIRK channel activity in the RGS7-KO was best reproduced with an approximately 90% decrease in RGS concentration. This suggests that a vast majority of dendritic RGS proteins in CA1 pyramidal cells are of the RGS7 type, however, the exact proportion may depend on whether the types of RGS of the remaining portion are more or less efficient in GTP hydrolysis and whether the spontaneous, non-RGS-catalyzed GTP hydrolysis also plays a significant role. The estimate may also be affected by the gradual decrease in the concentration of the GABABR agonist in the extracellular medium, which we lacked in our modelling framework. However, insights such as this can help in better characterising the RGS proteins’ contribution to GABABR-GIRK coupling, which is made obscure by the multitude of RGS subtypes and their interacting proteins [Doupnik, 2015]. Our model can be equally well used to complement the data on how presynaptic depression is affected by RGS proteins, reviewed in [Gerber et al., 2016], as well as other GABABR-pathway proteins.

The level of biochemical detail in our model makes it a strong candidate for dissecting the effects of GABABR activation on brain activity at multiple levels. Specifically, only neuron models in which the effects of GABABR activation are constrained by physiological limits — e.g., the strength of GABABR-mediated hyperpolarization at the postsynaptic dendrite should not exceed the maximal GIRK channel conductance — allow computational modeling to extract reliable information about the processes underlying the observed neuronal phenomena. In this work, we used our model to analyze data from [Palmer et al., 2012] (Fig. 4), shedding light on the neural mechanisms of interhemispheric inhibition of rodent L5PCs. Most of the characteristics of the firing in response to contralateral paw stimulation and its GABABR-mediated inhibition by ipsilateral paw stimulation were reproduced in our single-L5PC model. However, there was a mismatch between the sensitivity of the GABABR-mediated inhibition to the timing of the inputs (Fig. 4J,N). This mismatch could be due to the dynamic structure of the predicted L5PC firing pattern, which features an initial high-frequency firing phase (Fig. 4E–F), but it could also result from an unrealistic GABA input profile. Here, we explored the latter scenario, and found that our simulation results closely match experimental data on the timing sensitivity of GABA inputs when a ramp-down, rather than a square-pulse, form of GABABR activation was used (Figs. 4K and S2). This suggests that GABABR activation associated with interhemispheric inhibition may be strong shortly after the ipsilater paw stimulation and weaker later on. Such a scenario could be mediated by short-term depression either in the input [Price et al., 2005] or output [Oláh et al., 2009, Chittajallu et al., 2013, Abs et al., 2018] of the neurogliaform neurons, which are the main mediators of postsynaptic GABABR activation in the cortex [Oláh et al., 2009, Schulz et al., 2021]. Although the exact pattern of excitatory synaptic inputs to neurogliaform neurons during ipsilateral hindpaw stimulation is not known, their late spiking charateristics [Oláh et al., 2007] and the abovementioned observations of their synaptic depression make it plausible that the GABA signal activating the postsynaptic GABABRs was mediated by few APs in presynaptic neurogliaform neurons and had its strongest impact at the start of the spiking response of these neurons. This prediction, together with our analyses employing maximally reluctant presynaptic VGCCs (Fig. S3) or presynaptic VGCCs naturally inhibited through GABABR activation provide useful quantitative insights into the dynamics of the GABABR-mediated inhibition of L5PCs and the extent to which pre- and postsynaptic GABABR activation can inhibit L5PC firing in vivo.

The impact of GABABR activation at the tissue level is complex as it mediates postsynaptic GIRK channel activation and presynaptic short-term depression in both excitatory and inhibitory presynaptic terminals. Here, we modelled the effect of GABABR activation in all three phenomena, and we also compared the effects of pre- and postsynaptic GABABR activation on L5PC spiking under an in vivo-like synaptic stimulation. Future work should integrate all three modalities (contribution of GABABR activation to postsynaptic inhibitory currents, depression of E*→*E synapses, and depression of I*→*E synapses) into biophysically detailed recurrent network models to provide a more complete model on how manipulations of the GABABR system affect the total network excitability and oscillations. Importantly, when implemented together with the vesicle release model as in the experiments of Figs. 2 and 4, the model allows the analysis of how two types of presynaptic short-term depression, namely, the one mediated by GABABR activation and VGCC inactivation and the one mediated by depletion of readily releasable neurotransmitter vesicles, interact and counteract. This, together with future description of GABABR-meditated effects on the vesicular release machinery [Manz et al., 2019] and alternative mechanisms of short-term plasticity (such as one mediated by metabotropic glutamate receptors [Scanziani et al., 1997] or adenosine receptors [Brager and Thompson, 2003]), will allow a biophysically and biochemically detailed analysis that captures the most relevant molecular aspects of short-term dynamics in synapses. Although not considered here, the model can also be used for studying GABABR-activated GIRK currents in the presynaptic terminals, which have been observed in the cortex [Ladera et al., 2008].

GABABR activation inhibits voltage-gated Ca^2+^ channels also in the dendrites [Pérez-Garci et al., 2006, Palmer et al., 2012, Pérez-Garci et al., 2013, Schulz et al., 2021]. In this work, we only considered the presynaptic N-type Ca^2+^ channels, which display a strong shift in voltage-dependency in response to binding with G_i*βγ*_ [Colecraft et al., 2000], but notable effects have also been measured in P/Q-type channels [Huynh et al., 2015] which are also expressed presynaptically in many cell types. Previous work on non-neuronal cells also clearly shows the modulation of L-type channels by G_i*βγ*_ binding [Ivanina et al., 2000], but detailed information on how the modulation affects the conductance-voltage relationship is, to the best of our knowledge, missing, which hinders biophysically detailed modelling of these actions in neurons. However, the significance of GABABR-mediated inhibition of L-type Ca^2+^ channels in modulating pyramidal neuron firing activity has been shown [Pérez-Garci et al., 2013, Schulz et al., 2021], and the reports of N- and P/Q-type Ca^2+^ channels being expressed at pyramidal cell dendrites [Lorenzon and Foehring, 1995, Almog and Korngreen, 2014] suggest that the total contribution of VGCCs to the postsynaptic effects of GABABR activation may be substantial. In the modelling experiments of Fig. 3, we assumed that all inhibition of the L5PC firing by GABABR activation was mediated by GIRK channels. The experiments of [Schulz et al., 2021] showed that there was a significant interaction between nimodipine (an L-type Ca^2+^ channel inhibitor) application and stimulus intensity in modulating the effect of baclofen on spiking rate but that the nimodipine did not have a significant main effect. Likewise, in the modelling experiments of Fig. 4, we achieved a GABABR-mediated decrease in L5PC firing that was mediated by GIRK channel activation only, although the experiments of [Palmer et al., 2012] showed that some of the effects of baclofen on L5PC firing were mediated by VGCCs. Some of the estimates of the system-level parameters fitted in the experiments of Fig. 3 and 4 should therefore be revised once a functional model taking into account the effects of GABABR activation on dendritic VGCCs becomes available.

The GABABR system is a prominent candidate for the development of treatments for mental disorders [Evenseth et al., 2020]. Our model can be readily implemented in different brain regions, such as the striatum [Kupferschmidt and Lovinger, 2015], and used to study the effects of genetic disposition and pharmacological manipulations of the proteins involved in the GABABR network. In addition to manipulations of postsynaptic GABABR pathway proteins showcased here, our model can be used for biochemically detailed simulation of the constituents of presynaptic short-term depression, alterations of which have been postulated as a mechanism for addiction [Johnson and Lovinger, 2016]. Computational analyses of relevant neuron populations by our model could thus reveal mechanisms of many mental disorders and suggest novel treatment options.

## 5 Acknowledgements

We thank Andrei Kramer for assisting with the biochemical model conversion into SBML format via the SBtabVFGEN tool. Funding: Academy of Finland (318879, 330776, 336376, 355256, 358049), University of Oslo Convergence Environment (4MENT), Swedish Research Council (VR-M-2020-01652), European Union’s Research and Innovation Program Horizon Europe under grant agreement 101137289 (the Virtual Brain Twin Project) and 101147319 (EBRAINS 2.0 Project). The authors wish to acknowledge CSC Finland (project 2003397) and Sigma2 Norway (project NN9529K/NS9529K) for computational resources.

## Supplementary figures and tables

**Figure S1:**
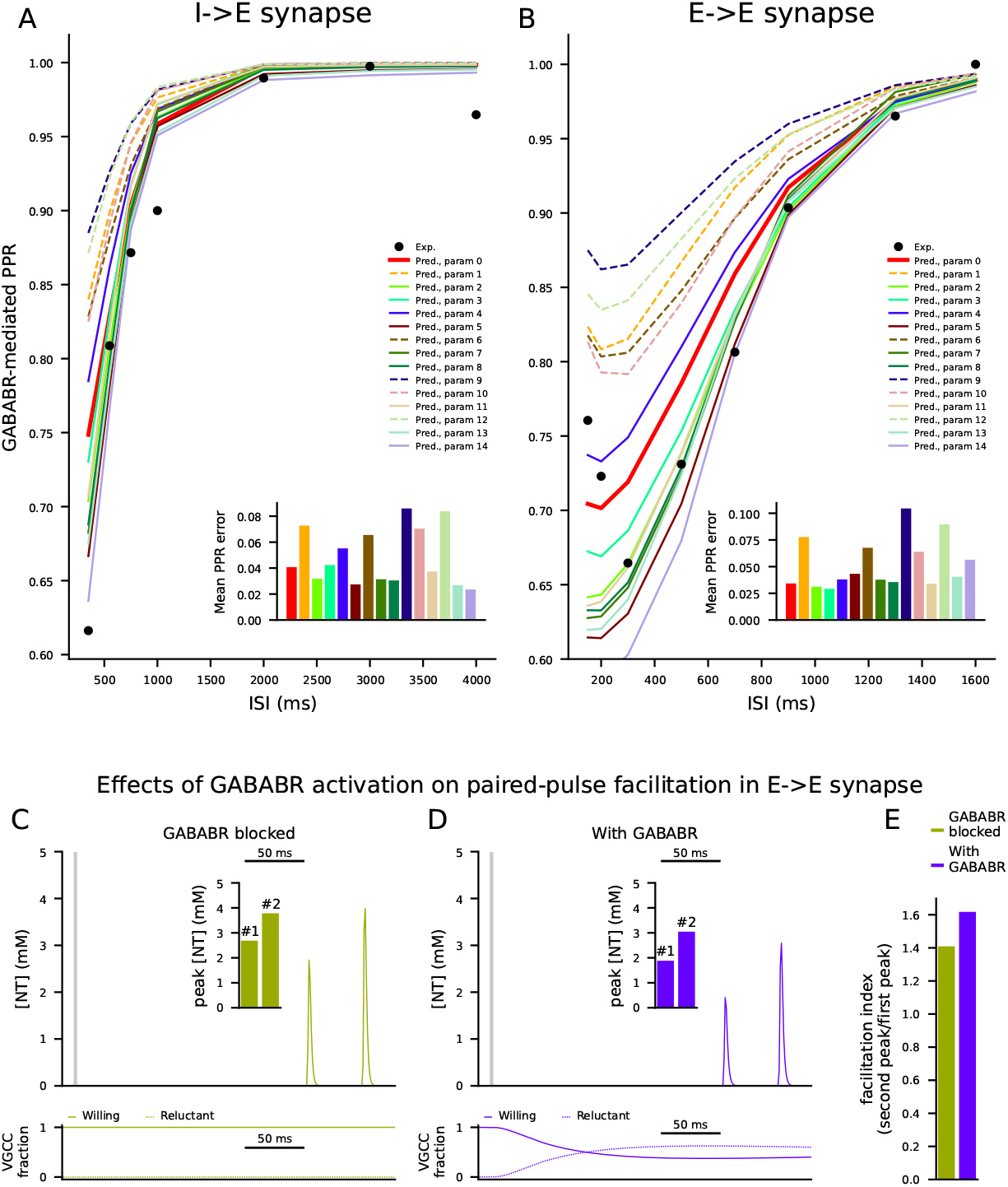
Supplementary results from the model with GABABRs at the presynaptic terminal. **A–B**: Most of the 15 models with best-fit parameters reproduce the PPR data also in the extended presynaptic model. The experiments of Fig. 2G,M were repeated using one of the 15 models with the best-fit parameters. The red, thick curve shows the simulation results for the default parameter set (see Fig. 2G,M). The dashed curves represent parameter sets for which the PPR prediction was off by more than 0.05 on average. **C–E**: The model reproduces a GABABR-dependency observed for paired-pulse facilitation of excitatory postsynaptic currents. The experiments of Fig. 2H–K were modified to model the paired-pulse facilitation protocol of [Isaacson et al., 1993]. Namely, the GABABR activating stimulus is given at time *t*=0 ms, and the first AP-inducing stimulation of the axon at *t*=200 ms, and the second AP-inducing stimulation at *t*=250 ms. C–D: The y-axes of the top panels show the glutamate concentration in the synaptic cleft when GABABRs were blocked (C) or functional (D), and the y-axes of the bottom panels show the fraction of VGCCs that were non-G_i*βγ*_-bound (solid) and G_i*βγ*_-bound (dashed). Inset: peak glutamate concentration following the first and the second axonal stimuli. These amplitudes were larger in the GABABR-active case (D) than in the GABABR-blocked case (C) because of the GABA-input-induced increase in the fraction of G_i*βγ*_-bound VGCCs. E: The facilitation index, calculated as the peak glutamate concentration following the second axonal stimulus divided by that following the first axonal stimulus, was larger in the GABABR-active than in the GABABR-blocked case, similar to experimental data [Isaacson et al., 1993].

**Figure S2:**
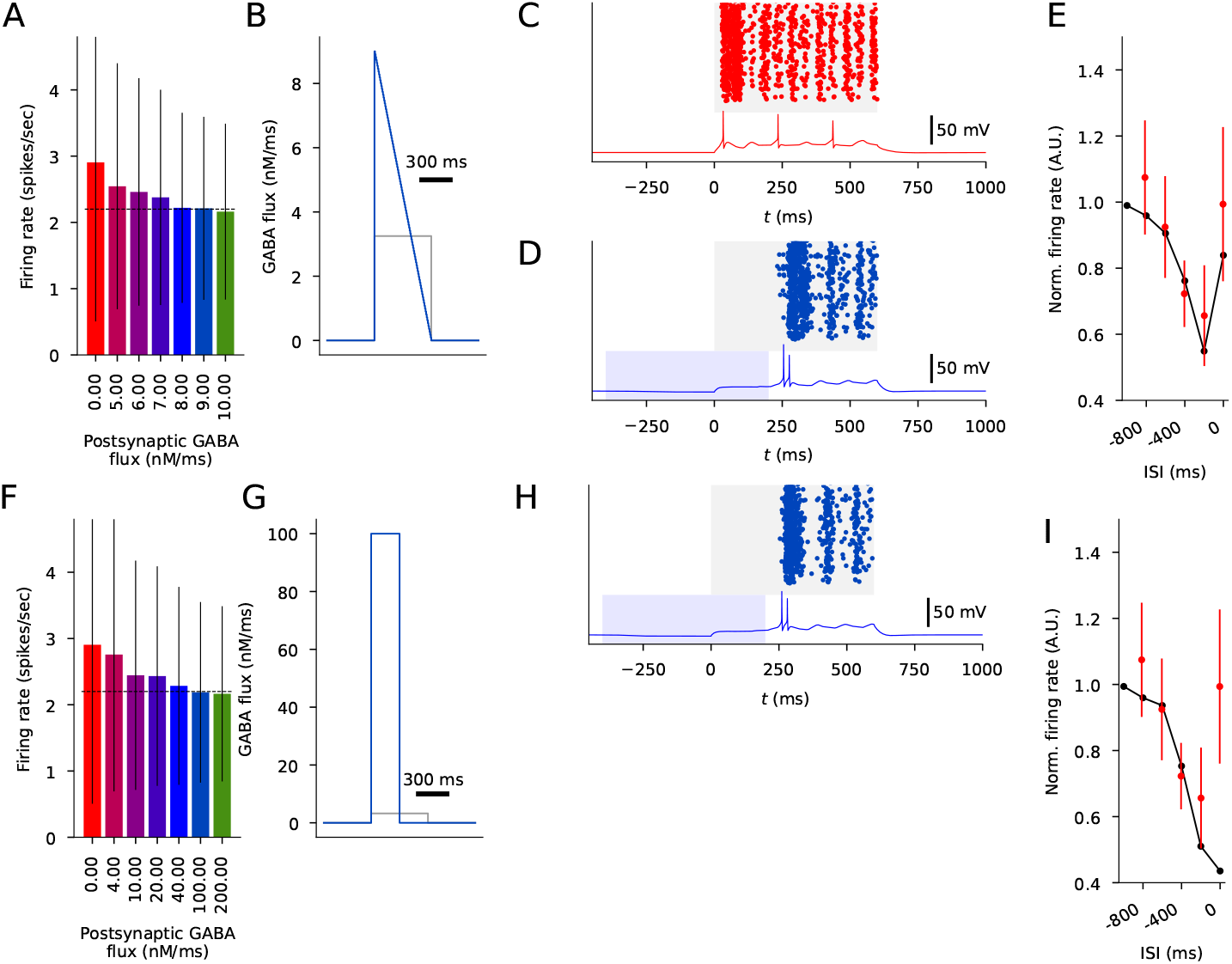
A ramp-down form but not a shorter square-pulse form of GABA input gives the model of interhemispheric inhibition the sensitivity to GABA timing that corresponds to experimental data. **A–E**: The simulations of Fig. 4 repeated using a ramp-down form of GABA input. **F–I**: The simulations of Fig. 4 repeated using a shorter (300 ms) square-pulse form of GABA input. **A**: The firing rate (spikes/sec) following the stimulus onset in the absence (red) and presence (other colors) of postsynaptic GABABR-activating inputs given different amplitudes of the ramp-down GABA flux. The onset of the GABA input was 400 ms before that of the glutamatergic inputs and lasted for 600 ms. The data of [Palmer et al., 2012] was best reproduced by using the GABA flux with a peak value 9 nM/ms. **B**: The form and amplitude of the GABA input used in Fig. 4 (gray) compared to the ramp-down form (blue). **C–D**: Illustration of the L5PC spiking across all trials and membrane potential time course of a single trial during the high-activity period (starting at *t* = 0 ms) in the absence (C) and presence (D) of the ramp-down GABA flux with a peak amplitude of 9 nM/ms. **E**: The normalized firing rate following the stimulus onset in the presence of postsynaptic GABABR-activating inputs with varied onset. The firing rates were normalized by the corresponding firing rate in the absence of GABABR-activating inputs. The red dots show the data measured in [Palmer et al., 2012], and the black curves show the average predictions of the model using the ramp-down GABA flux. The simulated firing rates are within 1 SD from the measured firing rates. **F**: The firing rate (spikes/sec) following the stimulus onset in the absence (red) and presence (other colors) of postsynaptic GABABR-activating inputs given different amplitudes of the shorter (300 ms) GABA flux. The data of [Palmer et al., 2012] was best reproduced by using the GABA flux of 100 nM/ms. **G**: The form and amplitude of the GABA input used in Fig. 4 (gray) compared to the shorter form (blue). **H**: Illustration of the L5PC spiking across all trials and a membrane potential time course in the presence of the 300-ms GABA flux with amplitude 100 nM/ms. **I**: The normalized firing rate following the stimulus onset in the presence of postsynaptic GABABR-activating inputs with varied onset. The simulations show a strong suppression for synchronous GABA and glutamatergic inputs (ISI = 0 ms), which is inconsistent with the experimental data.

**Figure S3:**
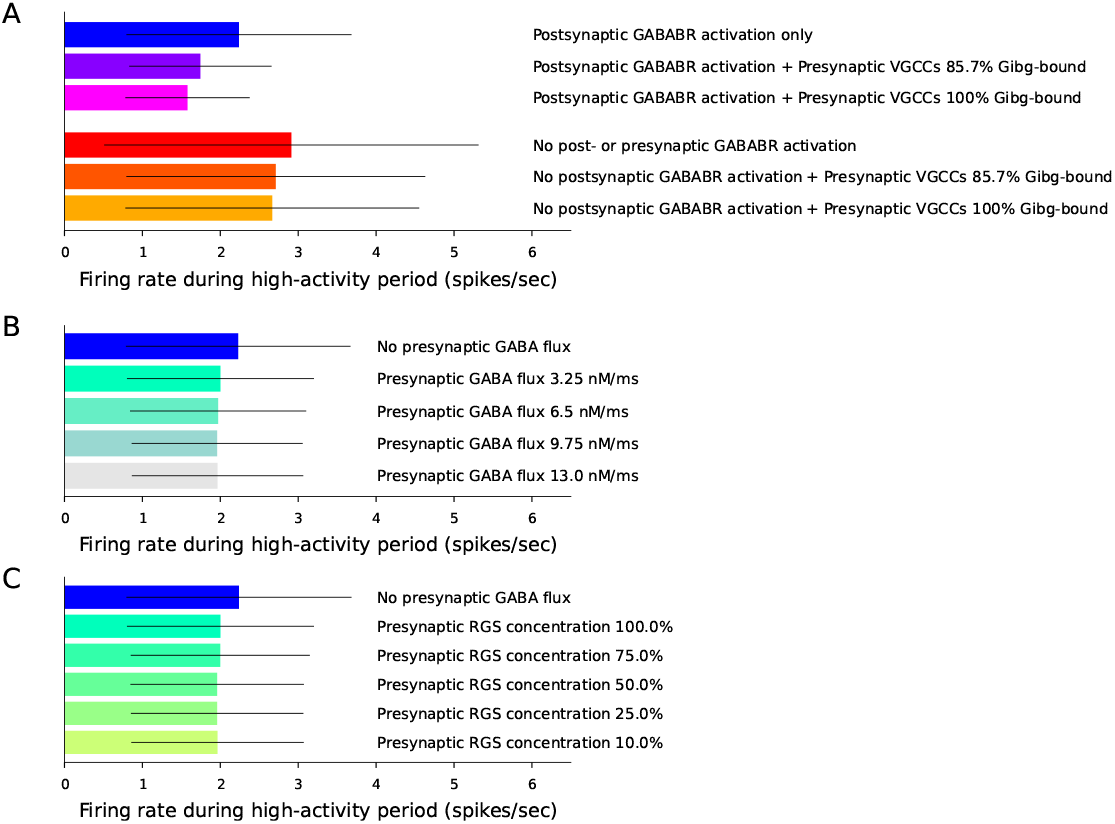
Exploration of the extent to which presynaptic GABABR activation can inhibit the L5PC firing during the high-activity period in the protocol of [Palmer et al., 2012]. **A**: Firing rates during the high-activity period (see Fig. 4) given the presence (top bars) or absence (bottom bars) of postsynaptic effects of GABABR activation and an external control over presynaptic VGCC inhibition. The three different bars in both categories show the firing rates with different externally set levels of G_i*βγ*_-bound VGCCs in the presynaptic terminal, namely, 0% (all willing), 85.7% (quasi-maximally reluctant), and 100% (maximally reluctant). **B**: Firing rates during the high-activity period given naturally occurring presynaptic GABABR activation assuming different amplitudes of GABA flux. The flux 3.25 nM/ms corresponds to the GABA flux given to the postsynaptic dendrites. **C**: Firing rates during the high-activity period given naturally occurring presynaptic GABABR activation assuming a flux of flux 3.25 nM/ms and different concentrations of presynaptic RGS proteins.

**Table S1:** Best-fit parameters from all 15 fitting tasks. *k*_1_, *k*_2_, *k*_3_, *k*_4_, and *k*_5_ are the reaction-rate factors for GABA extrusion (*k*_1_), GABA binding to GABABR (*k*_2_), overall GABABR dynamics (*k*_3_), and G_i*βγ*_ binding to (*k*_4_) and dissociation from (*k*_5_) GIRK and VGCC channels, see Table 2. The best-fit parameters used as the default parameter set in the result figures bolded.

| $k_1$ | $k_2$ | $k_3$ | $k_4$ | $k_5$ | [RGS<sub>post</sub>] | [RGS<sub>pre,E</sub>] | [RGS<sub>pre,I</sub>] | $f_{\text{GABA}}$ |
| --- | --- | --- | --- | --- | --- | --- | --- | --- |
| **107.034** | **514.651** | **167.633** | **11.026** | **66.931** | **3.009** $\mu\text{M}$ | **1.904** $\mu\text{M}$ | **2.241** $\mu\text{M}$ | **1507.740** particles/ms |
| 91.805 | 846.811 | 67.856 | 9.858 | 71.217 | 5.000 $\mu\text{M}$ | 1.817 $\mu\text{M}$ | 2.170 $\mu\text{M}$ | 3000.000 particles/ms |
| 133.092 | 528.522 | 185.581 | 8.574 | 25.868 | 3.186 $\mu\text{M}$ | 2.125 $\mu\text{M}$ | 2.729 $\mu\text{M}$ | 2815.426 particles/ms |
| 108.557 | 729.183 | 292.710 | 2.473 | 16.923 | 4.406 $\mu\text{M}$ | 2.127 $\mu\text{M}$ | 2.973 $\mu\text{M}$ | 696.729 particles/ms |
| 111.531 | 708.687 | 165.616 | 10.797 | 93.218 | 2.903 $\mu\text{M}$ | 1.728 $\mu\text{M}$ | 2.235 $\mu\text{M}$ | 2797.567 particles/ms |
| 109.869 | 469.365 | 114.137 | 59.053 | 73.263 | 3.573 $\mu\text{M}$ | 2.385 $\mu\text{M}$ | 3.028 $\mu\text{M}$ | 2852.993 particles/ms |
| 84.387 | 57.393 | 126.013 | 4.253 | 57.544 | 5.000 $\mu\text{M}$ | 1.722 $\mu\text{M}$ | 2.122 $\mu\text{M}$ | 929.204 particles/ms |
| 126.993 | 135.047 | 348.945 | 3.733 | 18.257 | 3.800 $\mu\text{M}$ | 2.647 $\mu\text{M}$ | 3.237 $\mu\text{M}$ | 2804.842 particles/ms |
| 105.438 | 642.913 | 142.107 | 19.602 | 40.482 | 3.599 $\mu\text{M}$ | 2.391 $\mu\text{M}$ | 2.974 $\mu\text{M}$ | 1386.488 particles/ms |
| 91.403 | 459.570 | 97.658 | 3.443 | 65.833 | 5.000 $\mu\text{M}$ | 1.540 $\mu\text{M}$ | 1.942 $\mu\text{M}$ | 2447.114 particles/ms |
| 71.410 | 380.140 | 143.521 | 2.991 | 45.796 | 10.000 $\mu\text{M}$ | 2.044 $\mu\text{M}$ | 2.836 $\mu\text{M}$ | 2126.204 particles/ms |
| 146.055 | 702.846 | 310.725 | 3.041 | 13.076 | 3.334 $\mu\text{M}$ | 2.351 $\mu\text{M}$ | 3.166 $\mu\text{M}$ | 2112.405 particles/ms |
| 90.417 | 993.718 | 98.403 | 6.879 | 95.287 | 3.944 $\mu\text{M}$ | 1.572 $\mu\text{M}$ | 2.204 $\mu\text{M}$ | 1413.654 particles/ms |
| 119.681 | 591.014 | 91.960 | 89.687 | 88.948 | 3.453 $\mu\text{M}$ | 2.300 $\mu\text{M}$ | 2.755 $\mu\text{M}$ | 2950.763 particles/ms |
| 123.170 | 616.560 | 117.913 | 74.842 | 46.625 | 3.952 $\mu\text{M}$ | 2.609 $\mu\text{M}$ | 3.283 $\mu\text{M}$ | 3000.000 particles/ms |

## Notes

### Competing Interest Statement

Jan M. Schulz works as a senior scientist at Roche. Ole A. Andreassen has received speaker fees from Lundbeck, Janssen, Otsuka, and Sunovion and is a consultant to Cortechs.ai and Precision Health. These affiliations did not influence the conduct of the study or the preparation of this manuscript.

